# Delineating regional vulnerability in the neurodegenerative disease SCA1 using a conditional mutant ATXN1 mouse

**DOI:** 10.1101/2023.02.08.527710

**Authors:** Lisa Duvick, W. Michael Southern, Kellie Benzow, Zoe N. Burch, Hillary P. Handler, Jason S. Mitchell, Hannah Kuivinen, Udaya Keerthy Gadiparthi, Praseuth Yang, Alyssa Soles, Carrie Scheeler, Orion Rainwater, Shannah Serres, Erin Lind, Tessa Nichols-Meade, Brennon O’Callaghan, Huda Y. Zoghbi, Marija Cvetanovic, Vanessa C. Wheeler, James M. Ervasti, Michael D. Koob, Harry T. Orr

## Abstract

Spinocerebellar ataxia type 1 (SCA1) is a fatal neurodegenerative disease caused by an expanded polyglutamine tract in the widely expressed ATXN1 protein. To elucidate anatomical regions and cell types that underlie mutant ATXN1-induced disease phenotypes, we developed a floxed conditional knockout mouse model (*f-ATXN1^146Q/2Q^*) having mouse *Atxn1* coding exons replaced by human exons encoding 146 glutamines. *F-ATXN1^146Q/2Q^* mice manifest SCA1-like phenotypes including motor and cognitive deficits, wasting, and decreased survival. CNS contributions to disease were revealed using *ATXN1^146Q/2Q^*;*Nestin-Cre* mice, that showed improved rotarod, open field and Barnes maze performances. Striatal contributions to motor deficits were examined using *f-ATXN1^146Q/2Q^*;*Rgs9-Cre* mice. Mice lacking striatal *ATXN1^146Q/2Q^* had improved rotarod performance late in disease. Muscle contributions to disease were revealed in *f-ATXN1^146Q/2Q^*;*ACTA1-Cre* mice which lacked muscle pathology and kyphosis seen in *f-ATXN1^146Q/2Q^* mice. Kyphosis was not improved in *f-ATXN1^146Q/2Q^;Nestin_-_Cre* mice. Thus, optimal SCA1 therapeutics will require targeting mutant ATXN1 toxic actions in multiple brain regions and muscle.

## INTRODUCTION

Neurodegenerative diseases are often characterized by a variety of clinical phenotypes. Thus, two fundamental issues facing neurodegenerative disease research are 1) the identification of the anatomical/cellular features underlying each disease symptom, and 2) assessing the extent to which disease processes in these regions are similar at a molecular level. Progress in these areas provides vital information for developing treatments to ensure maximum therapeutic efficacy. In addition, mapping the anatomical basis of disease-associated phenotypes/symptoms is a means of further understanding CNS function at a systems level.

Spinocerebellar ataxia type 1 (SCA1), an autosomal dominant neurodegenerative disease, is one of the nine neurodegenerative diseases caused by expansion of a CAG trinucleotide tract encoding a polyglutamine stretch in the affected ataxin-1 (ATXN1) protein^1,2^. The initial symptom of SCA1 is typically gait ataxia. As the disease progresses, common symptoms include dysarthria, muscle wasting, and cognitive deficits such as executive functioning difficulties. Late-stage disease involves bulbar dysfunction, which is thought to cause the swallowing and breathing difficulties that underlie premature death^3-7^. The typical pathological pattern is olivopontocerebellar atrophy with loss of cerebellar Purkinje cells, major neuronal loss in the dentate nuclei, and extensive olivary neuronal loss. Basal pontine neuronal atrophy is variable and, in some cases, severe. Other brainstem areas affected are the red nuclei, the vestibular nuclei, and motor cranial nerve nuclei^4,7^. In the spinal cord, the anterior horns, posterior columns, and spinocerebellar tracts also have atrophy with variable sparing of the pyramidal tracts. The pars compacta of the substantia nigra is relatively spared, but the pallidum and thalamus can have mild involvement. The basal forebrain cholinergic nuclei, cerebral cortex, and hippocampus may have mild neuronal loss^4,7^. Previous studies demonstrate the utility of *Atxn1* knockin mice, in which an expanded stretch of CAG repeats is inserted into one *Atxn1* allele, as a model for studying the spectrum of SCA1-like phenotypes^8^. In this study, we describe the generation and characterization of a conditional SCA1 knockin mouse model in which the two murine *Atxn1* coding exons, 7 and 8, in one allele were replaced with the human *ATXN1* coding exon 8, containing an expanded 146 CAG repeat, and human exon 9. LoxN recombination sites flank the human *ATXN1* exons such that the human coding exons of the expanded *ATXN1^146Q/2Q^* allele are deleted in the presence of Cre recombinase. Heterozygous floxed humanized *ATXN1^146Q/2Q^* (*f-ATXN1^146Q/2Q^*) mice manifest a very similar collection of SCA1-like phenotypes as seen in *Atxn1^154Q^* knockin mice^8^, i.e. a progressive neurological disorder featuring motor incoordination, cognitive deficits, wasting with kyphosis, and decreased survival. *F-ATXN1^146Q/2Q^* mice were crossed with *Nestin-Cre* and *Rgs9*-*Cre* mice to reveal contributions of all CNS cells and striatal medium spiny neurons (MSNs) to SCA1-like phenotypes, respectively. Further, *f-ATXN1^146Q/2Q^* were crossed to *ACTA1-Cre* mice, to assess the contribution of skeletal muscle to SCA1-like phenotypes. By biochemical and image analyses, *f*-*ATXN1^146Q/2Q^; ACTA1-Cre* mice showed a strong peripheral muscle specific phenotype in *f*-*ATXN1^146Q/2Q^* mice. The findings from studying each of these sub-populations involved in disease progression has important implications for the biological targeting of SCA1 therapeutic agents.

## RESULTS

### Generation and characterization of the conditional knockout *f-ATXN1^146Q/2Q^*SCA1 mouse model

A conditional knockout mouse model designated *f*-*ATXN1^146Q/2Q^* was developed to investigate brain region and peripheral tissue contributions to SCA1-like phenotypes as well as to obtain a platform for testing human *ATXN1* gene targeting therapies. The entire coding region of one allele of the mouse *Atxn1* gene was replaced with that of the human *ATXN1* gene in a two-step process. First, the 3’ end of the *Atxn1* gene (20kb) that includes the last two exons through the end of the transcript (Figure1ai) was deleted from the genome of a mouse ES cell line (C57BL/6N) and replaced by Frt recombination sites and selectable markers (Figure1aii). Next, the syntenic sequence from the human genome (31kb) was inserted using site-specific recombination at flanking FRT sites (Figure1aiii). The ES cell was injected into C57Bl6 embryos yielding *f-ATXN1^146Q/^*^-^ mice, which were then crossed with WT mice to yield *f-ATXN1^146Q/2Q^* mice (Figure 1aiv). With respect to coding sequences, these mice have one mouse WT allele and one human allele encoding ATXN1 containing 146 CAG repeats. In this study, *f-ATXN1^146Q/2Q^* mice were crossed with various Cre recombinase expressing mice to delete the *f-ATXN1^146Q^* allele from select regions/tissues.

**Figure 1.**
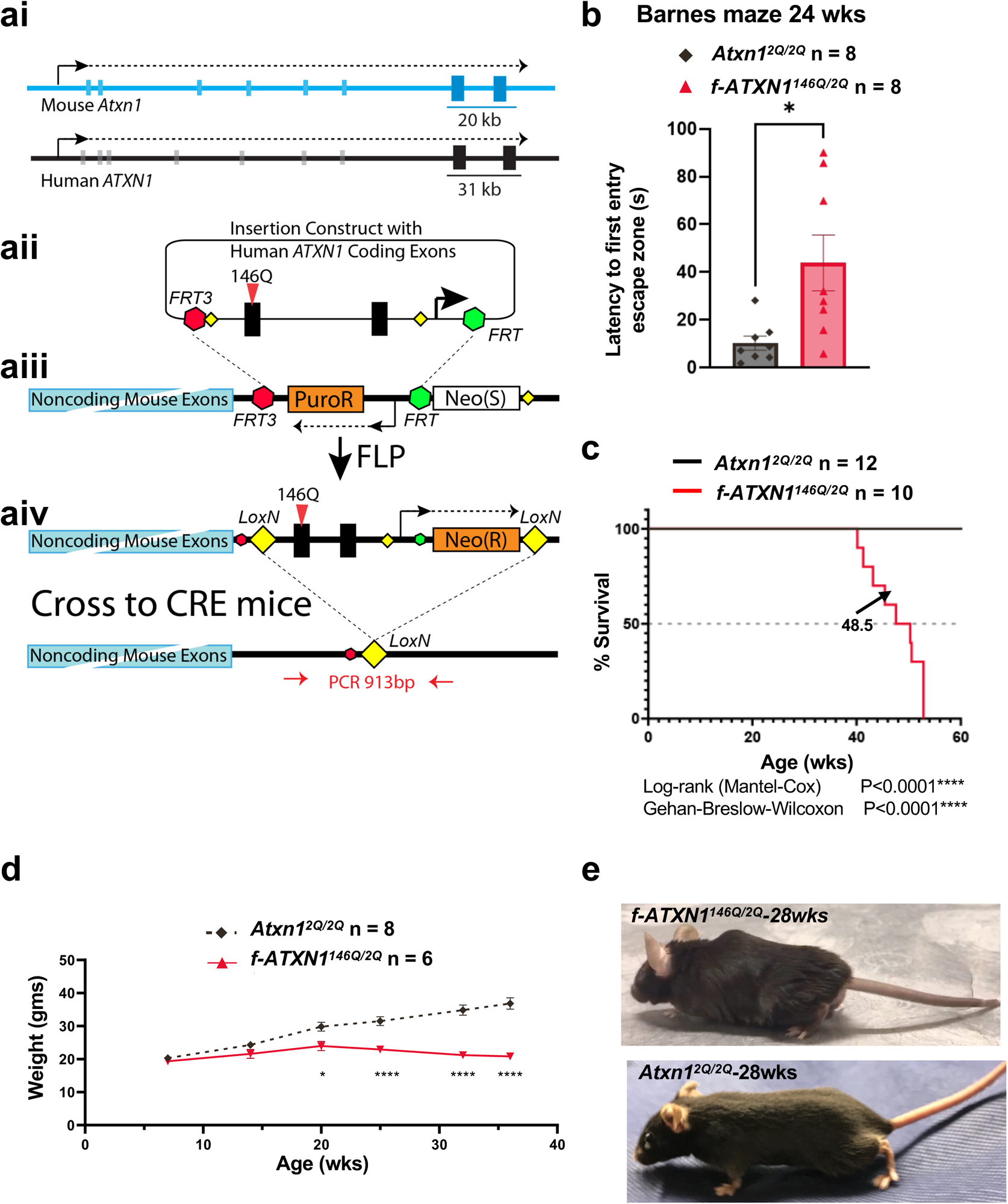
Generation and characterization of the *f-Atxn1^146Q/2Q^* conditional mouse model. **ai**, Organization of the mouse *Atxn1* (blue) and human *ATXN1* (black) genes, with the only two exons encoding the ATXN1 protein indicated by boxes larger and darker than the non-coding exons. The size (kb) and location of the mouse genomic sequences (blue) replaced by the human genomic sequences (black) in the *f-Atxn1^146Q^* allele are indicated. **ii,** The portion of the *Atxn1* gene encompassing the two coding exons was replaced with an *FRT*-recombination recipient cassette in mouse ES cells and then that cassette was replaced with the portion of the human *ATXN1* genomic sequences syntenic to the deleted mouse sequence using FLP recombinase. **iii,** The inserted human sequences in the resulting *Atxn1^146Q^* allele are flanked by *LOX* recombination sites, as shown. **iv,** Mating mice with this allele to lines expressing CRE recombinase removes the human *ATXN1* insertion, as shown. **b**, Barnes Maze performance at 24 weeks of age, on-way ANOVA Tukey’s post hoc test. **c,** Mouse survival plotted as Kaplan-Meyer curves with median lifespan labeled for each genotype. **d**, Body weight measurements between 6 and 36 weeks of age, two-way ANOVA Bonferroni’s post hoc test. **e,** Representative photographs of 28-week-old *Atxn1^2Q/2Q^* and *f-ATXN1^146Q/2Q^* showing kyphosis. Significance of results is denoted as * (p<0.05), ** (p<0.01), *** (p<0.001), and **** (p<0.0001).

To evaluate the ability of the *ATXN1^146Q^* allele to be deleted by Cre recombinase, *f-ATXN1^146Q/2Q^* mice were crossed with *Sox2-Cre* mice^9^. This *Sox2-Cre* mouse mediates efficient Cre-mediated recombination at gastrulation in all epiblast-derived cells including the primordial germ cells. Using CAG repeat-specific and recombination-specific PCR primers (Figure S1a), recombination and deletion of the *ATXN1^146Q^* allele was detected in all tissues examined from a *f-ATXN1^146Q/2Q^; Sox-2-Cre* mouse (Figures. S1b-d).

Expression of the ATXN1[146Q] protein was assessed by immunoblotting protein extracts prepared from three brain regions from WT, *f-ATXN1^146Q/2Q^*, and *Atxn1^175Q/2Q^* mice, the latter derived from the *ATXN1^154Q/2Q^* knockin line developed by Watase et al.^8^ In two-week-old mice, the levels of ATXN1[146Q] and ATXN1[175Q] relative to mouse ATXN1[2Q] or TUBULIN were very similar in the three brain regions examined, cerebral cortex, cerebellum, and brainstem (Figures S2a-S2c). Expression of the *f-ATXN1^146Q^* transcript was also examined by qRT-PCR (Figure 2Sd). Total RNA was extracted from the cerebellum of two-week-old *f-ATXN1^146Q/2Q^* mice. As shown in Figure S2e human *f-ATXN1^146Q^* RNA levels are not statistically different from mouse *Atxn1^2Q^* RNA levels in the cerebellum.

*F-ATXN1^146Q/2Q^* mice display the spectrum of SCA1-like phenotypes observed in *Atxn1^154Q/2Q^* mice^8^. The SCA-like phenotypes exhibited by *f-ATXN1^146Q/2Q^* mice include a cognitive deficit on the Barnes Maze (Figure 1b), reduced survival (Figure 1c) and wasting (Figure 1d). In addition, Figure 1e shows that *f-ATXN1^146Q/2Q^* mice also have prominent kyphosis. Typical of SCA1 mouse models, *f-ATXN1^146Q/2Q^* mice have a motor performance deficit as assessed by accelerating rotarod assay (see below).

### Impact of deleting the *ATXN1^146Q^* allele from the CNS on SCA1-like phenotypes

Nestin-Cre mice express Cre recombinase in neuronal and glial precursors and thus throughout the CNS^10^. *F-ATXN1^146Q/2Q^; Nestin-Cre* mice were examined to assess the extent to which deletion of the *f-ATXN1^146Q^* allele from CNS neuronal and glial cells improved other SCA1-like neurological phenotypes manifested by *f-ATXN1^146Q^* mice. Figure 2a shows that *f-ATXN1^146Q/2Q^;Nestin-Cre* mice had a significant improvement in performance in the open-field test (locomotion) compared to *f-ATXN1^146Q/2Q^* mice at 12 weeks of age. *F-ATXN1^146Q/2Q^; Nestin-Cre* mice had improved cognitive performance as evaluated by the Barnes Maze test. Compared to controls *Atxn1^2Q/2Q^* and *Atxn1^2Q/2Q^; Nestin-Cre* mice, *f-ATXN1^146Q/2Q^* mice showed a significant delay in their first entry to the escape hole zone while *f-ATXN1^146Q/2Q^; Nestin-Cre* mice did not (Figure 2b). *F-ATXN1^146Q/2Q^; Nestin-Cre* mice had significant improvement in survival (Figure 2c) and slightly less, but significant wasting compared to *f-ATXN1^146Q/2Q^* mice (Figure 1d). In addition, at 36 weeks of age *f-ATXN1^146Q/2Q^* displayed a hindlimb clasping phenotype characteristic of mice with pathology in a variety of brain regions (Figure 2e) ^11^. The hindlimb clasping phenotype was normal in *f-ATXN1^146Q/2Q^; Nestin-Cre* mice at this age.

**Figure 2.**
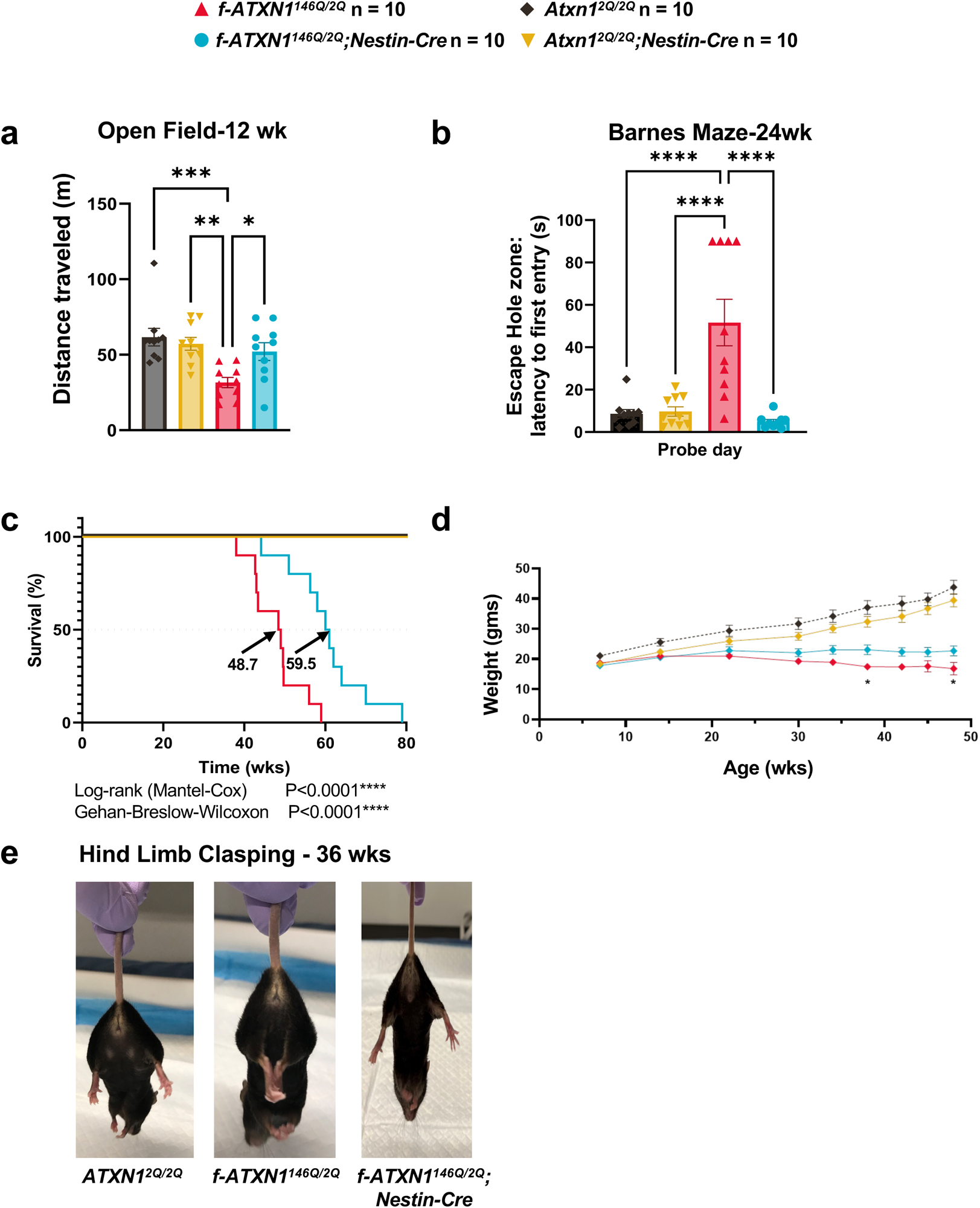
SCA1-like phenotypes improved in *f-ATXN1^146Q/2Q^*; *Nestin-Cre* mice. **a,** Open field performance at 12 weeks of age, one-way ANOVA Tukey’s post hoc test. **b**, Barnes Maze performance at 24 weeks of age, one-way ANOVA Tukey’s post hoc test. **c,** Mouse survival plotted as Kaplan-Meyer curves with median lifespan labeled for each genotype. **d**, Body weight measurements between 7 and 48 weeks-of-age. two-way ANOVA Bonferroni’s post hoc test. **e,** Hind limb clasping at 36 weeks. Significance of results is denoted as * (p<0.05), ** (p<0.01), *** (p<0.001) and ****(p<0.0001).

Naïve *f-ATXN1^146Q/2Q^* mice have a significant deficit in performance on the accelerating rotarod at 6 weeks, which progresses in severity with age (Figure 3a). Notably, *f-ATXN1^146Q/2Q^;Nestin-Cre* mice with the *ATXN1^146Q^* allele deleted throughout the CNS^10^ (Figures S3a, S3b & S3d), displayed improved rotarod performance at all ages tested, from 6 to 31 weeks-of-age (Figure 3b and Figure S4a). Previous studies in which the cerebella of 5 week old SCA1 knockin mice were injected with adeno-associated viruses expressing inhibitory RNAs targeting *Atxn1* rescued rotarod performance, indicating a role for the cerebellum in the early motor deficit seen in SCA1^12,13^. Of note, in SCA1 patients, changes in striatal volume are associated with an age-associated decline in motor performance^14,15^. Thus, as a first step in assessing regions of the CNS in addition to the cerebellum where dysfunction might contribute to altered rotarod performance, *f-ATXN1^146Q/2Q^* mice were crossed to *Rgs9-Cre* mice to delete the *ATXN1^146Q^* allele from striatal MSNs^16,17^ (Figure S5). Interestingly, deletion of *ATXN1^146Q^* from striatal MSNs yielded an improvement of rotarod performance that did not manifest until 31 weeks-of-age in *f-ATXN1^146Q/2Q^/Rgs9-Cre* mice relative to *f-ATXN1^146Q/2Q^*mice (Figure 3c and Figure S4b). Deletion of *ATXN1^146Q^* from striatal MSNs significantly improved dopamine- and cAMP-regulated phosphoprotein-32 (DARPP-32) expression in the striatum of *f-ATXN1^146Q/2Q^/Rgs9-Cre* mice (Figures S6a and S6b).

**Figure 3.**
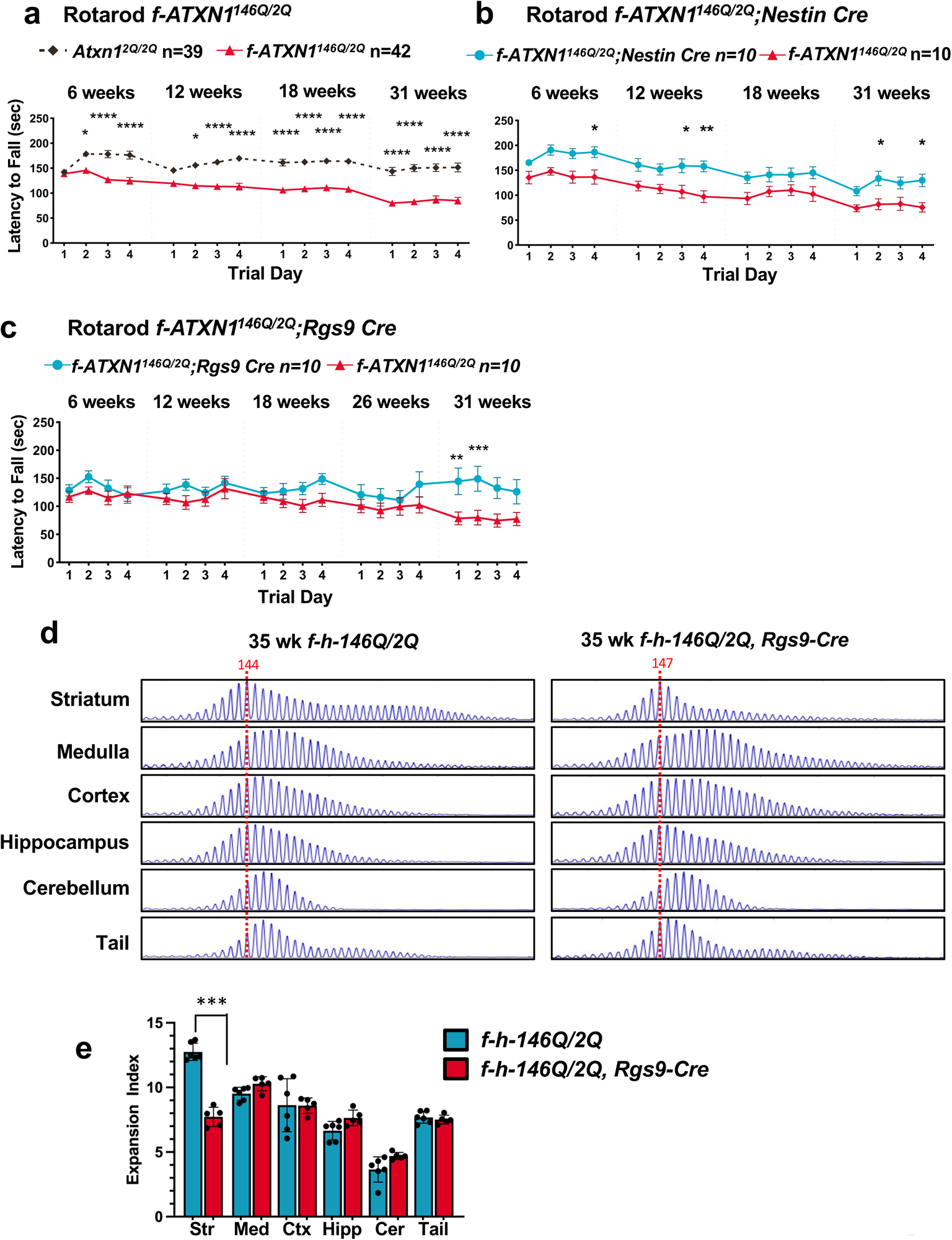
Striatal pathology contributes to the progressive motor performance deficit in *f-Atxn1^146Q/2Q^* mice. **a,** Rotarod assessment of combined data for *Atxn1^2Q/2Q^* and *f-ATXN1^146Q/2Q^* mice across 4 trial days at 6,12,18, and 31 weeks of age. **b,** Rotarod assessment of *f-Atxn1^146Q/2Q^* and *f-ATXN1^146Q/2Q^; Nestin-Cre.* **c,** Rotarod assessment of *f-Atxn1^146Q/2Q^* and *f-ATXN1^146Q/2Q^; Rgs9-Cre.* **a-c** Two-way repeated measures ANOVAs with Šídák post hoc test. Significance of results is denoted as * (p<0.05), ** (p<0.01), *** (p<0.001), and **** (p<0.0001). **d,** Representative Gene Mapper traces showing somatic instability in 35-week *f-Atxn1^146Q/2Q^* and *f-Atxn1^146Q/2^*; *Rgs9*-Cre mice. **e,** Quantified expansion indices. *f-Atxn1^146Q/2Q^* n=6 (3M, 3F), CAG 142-147 (determined from stable striatal peak = red dotted line); *f-Atxn1^146Q/2Q^*; *Rgs9*-Cre n=5 (3M, 2F), CAG 145-138. Paired t-test **** (p<0.0001). Small increases in expansion indices (p<0.05) in *f-Atxn1^146Q/2Q^*; *Rgs9*-Cre medulla, hippocampus and cerebellum may be due to slightly higher CAG lengths in these mice relative to *f-Atxn1^146Q/2Q^*.

Previous studies have shown extensive somatic expansion of the *ATXN1* CAG repeat in SCA1 patient and knock-in mice striatum^18,19^. Analyses of 35-week *f-ATXN1^146Q/2Q^*mice also revealed high levels of somatic expansion in striatum. Significant CAG expansion was also observed in other tissues/brain regions, most notably the medulla (Figures 3d and 3e), with *f-ATXN1^146Q/2Q^* mice exhibiting a brain regional expansion profile similar to that in a SCA1 patient^19^. Striatal expansion was significantly reduced in *f-ATXN1^146Q/2Q^/Rgs9-Cre* mice (Figures 3d and 3e), demonstrating that expansions occur in MSNs. Residual striatal expansion in *f-ATXN1^146Q/2Q^/Rgs9-Cre* mice may be explained by incomplete Cre-mediated excision of the expanded CAG repeat-containing allele in MSNs and/or CAG expansion in other striatal cell types (Figure S3a). We conclude that, as in SCA1 patients, disease in the striatum contributes to deficits in motor performance with disease progression in *f-ATXN1^146Q/2Q^* mice. Our results indicate that somatic CAG expansion in the striatum, and in particular in striatal MSNs may contribute to the development of this late motor deficit.

### Deletion of the *f-ATXN1*^146Q^ allele from skeletal muscle reveals a direct muscle-specific pathogenic effect of ATXN1[146Q]

In *mdx* mice, kyphosis is associated with paralumbar skeletal muscle atrophy^20^. Figure 4a shows that *f-ATXN1^146Q/2Q^* mice manifest a severe kyphosis by 20 wks with increasing severity to 50 weeks of age. We examined the CNS impact of ATXN1[146Q] on kyphosis by deleting ATXN1[146Q] from all CNS neurons, including spinal motor neurons by crossing *f-ATXN1^146Q/2Q^* mice with *Nestin-Cre* mice. Image analysis showed that kyphosis in *f-ATXN1^146Q/2Q^; Nestin-Cre* mice remained severe (Figure 4b). To examine whether kyphosis in *f-ATXN1^146Q/2Q^* mice is due to a direct effect of ATXN1[146Q] on muscle, *f-ATXN1^146Q/2Q^* mice were crossed with *ACTA1-Cre* mice to delete the *ATXN1^146Q^* allele from muscle^21^. Figures S3c and S3e show that in *f-ATXN1^146Q/2Q^; ACTA1-Cre* mice, expression of the *ATXN1^146Q^* allele was significantly reduced in skeletal muscle, including the tongue and diaphragm but not in the heart, cerebral cortex, cerebellum, medulla, or lung. *ACTA1-Cre*-induced deletion of *ATXN1^146Q^* eliminated the kyphosis seen in *f-ATXN1^146Q/2Q^* mice at 20 and 50 weeks of age (Figure 4c). Using the approach depicted in Figure 4e to calculate a kyphosis index (KI), the KI of *f-ATXN1^146Q/2Q^* mice is significantly reduced compared to the KI of *Atxn1^2Q/2Q^* mice (Figure 4f). Notably, deletion of the *ATXN1^146Q^* allele from muscle restored the KI to that of *Atxn1^2Q/2Q^* mice. These data support the concept that kyphosis in *f-ATXN1^146Q/2Q^* mice is a result of muscle atrophy induced by expression of ATXN1[146Q] in muscle.

**Figure 4.**
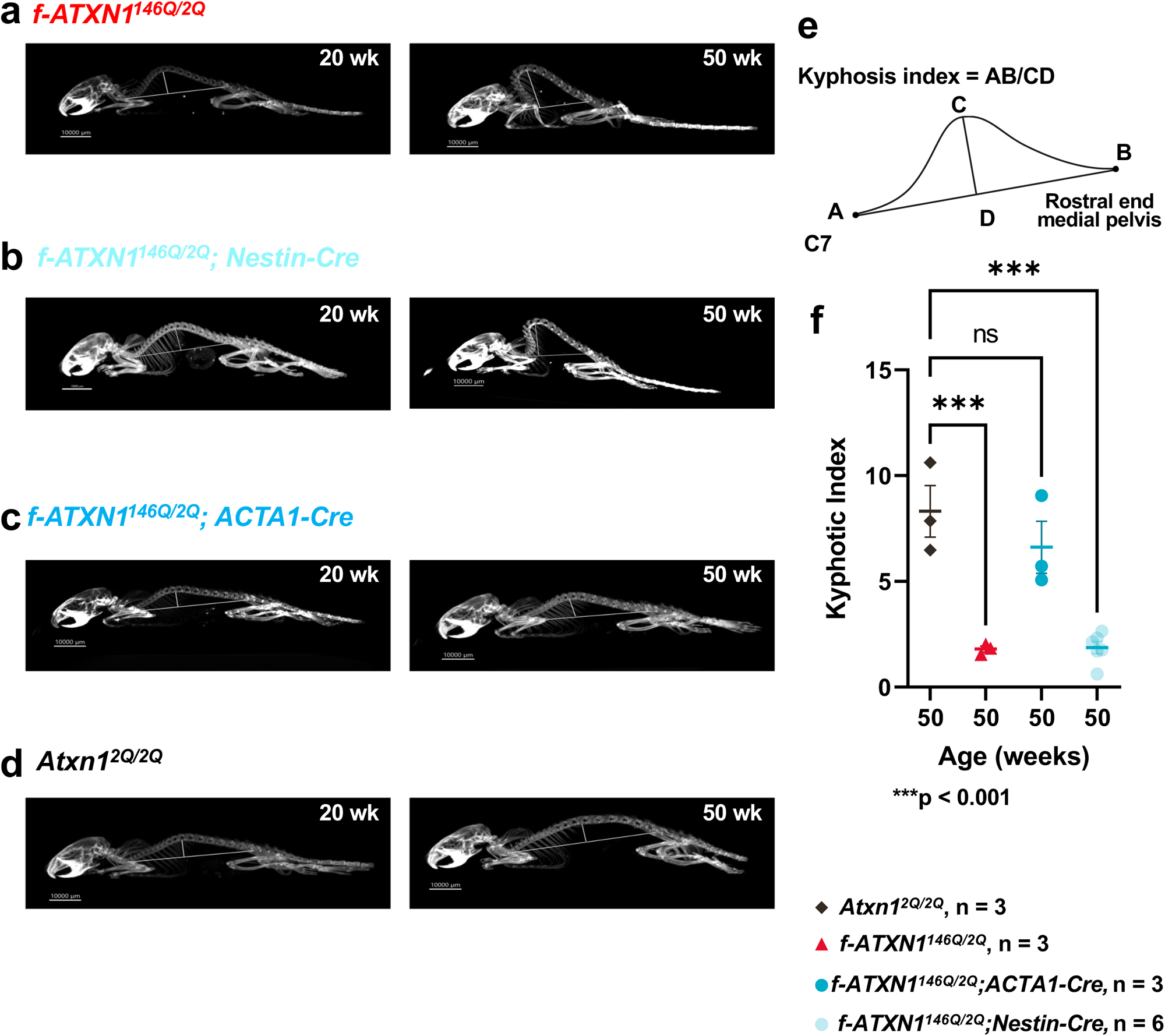
Progression of Kyphosis pathology in *f-ATXN1^146Q/2Q^* mice is corrected with deletion of *ATXN1^146Q^* in skeletal muscle. **a,** CT scan image of *f-ATXN1^146Q/2Q^* mice. **b,** CT scan image of *f-ATXN1^146Q/2Q^; Nestin-Cre* mice. **c,** CT scan image of *f-ATXN1^146Q/2Q^; ACTA1-Cre* mice. **d,** CT scan image of *Atxn1^2Q/2Q^*mice. **e-f,** Kyphosis image calculation and kyphosis index at 50 wks, one-way ANOVA with Dunnett’s post hoc test. Significance of results is denoted as * (p<0.05), ** (p<0.01), and *** (p<0.001).

Consistent with an ATXN1[146Q]-induced muscle atrophy phenotype, tibialis anterior (TA) and extensor digitorum longus (EDL) muscle masses were lower in *f-ATXN1^146Q/2Q^* mice compared to *Atxn1^2Q/2Q^*;*ACTA1-Cre* or *f-ATXN1^146Q/2Q^*;*ACTA1-Cre* mice at both 18 and 30 weeks (Figures 5a and 5h). Additional characterization of the effects of ATXN1[146Q] on skeletal muscle function revealed a progressive, muscle specific myopathy. Further evaluation of muscle strength was performed using an *in vivo* electrode-based assay that measures dorsiflexor muscle torque via direct stimulation of the peroneal nerve. Dorsiflexor (EDL + TA) torque was significantly lower at 18 wks and trending lower at 30 wks, in *f-ATXN1^146Q/2Q^* mice compared to *Atxn1^2Q/2Q^*;*ACTA1-Cre* mice, despite correcting for body mass or muscle mass (Figure 5b, 5c and 5i, 5j). To further investigate the ATXN1[146Q]-induced muscle weakness phenotype, we used an *ex vivo* assay that assesses EDL contractile function independent of endogenous motor neuron activation. At 18 wks, EDL specific force production was not different between genotypes suggesting that intrinsic muscle contractility, independent of motor neuronal activation, was not affected in *f-ATXN1^146Q/2Q^* mice (Figure 5d). However, at 30 wks, EDL specific force was lower in the *f-ATXN1^146Q/2Q^* mice, a phenotype that was fully corrected with deletion of the *ATXN1^146Q^* allele from muscle (Figure 5k). The *in vivo* but not *ex vivo* muscle strength deficits in *f-ATXN1^146Q/2Q^* mice at 18 weeks suggest the presence of a neuronal pathology potentially related to disrupted motor neuron activation or neuromuscular junction (NMJ) dysfunction. Consistent with possible alterations in NMJ function, at eight weeks of age increased expression of mRNAs encoding the NMJ proteins *MuSK* and *Chrna1*^22^ were detected (Figure S6c). At 40 weeks of age expression of *MuSK* and *Chrna1* RNAs were restored to normal levels in *Acta-Cre; f-ATXN1^146Q/2Q^* mice (Figure 6a) but not in *Nestin-Cre; f-ATXN1^146Q/2Q^*mice (Figure 6b). Furthermore, the reduction in EDL specific force observed at 30 weeks in *f-ATXN1^146Q/2Q^* mice could indicate a progressive myopathy stemming from a combination of chronic muscle disuse and alterations in intrinsic muscle contractile function. Importantly, all muscle deficits were restored to *Atxn1^2Q/2Q^;ACTA1-Cre* levels in mice with the *ACTA1-Cre* induced deletion of the *ATXN1^146Q^* allele. Together, these data indicate the presence of a progressive myopathy that eventually affects intrinsic muscle contractility.

**Figure 5.**
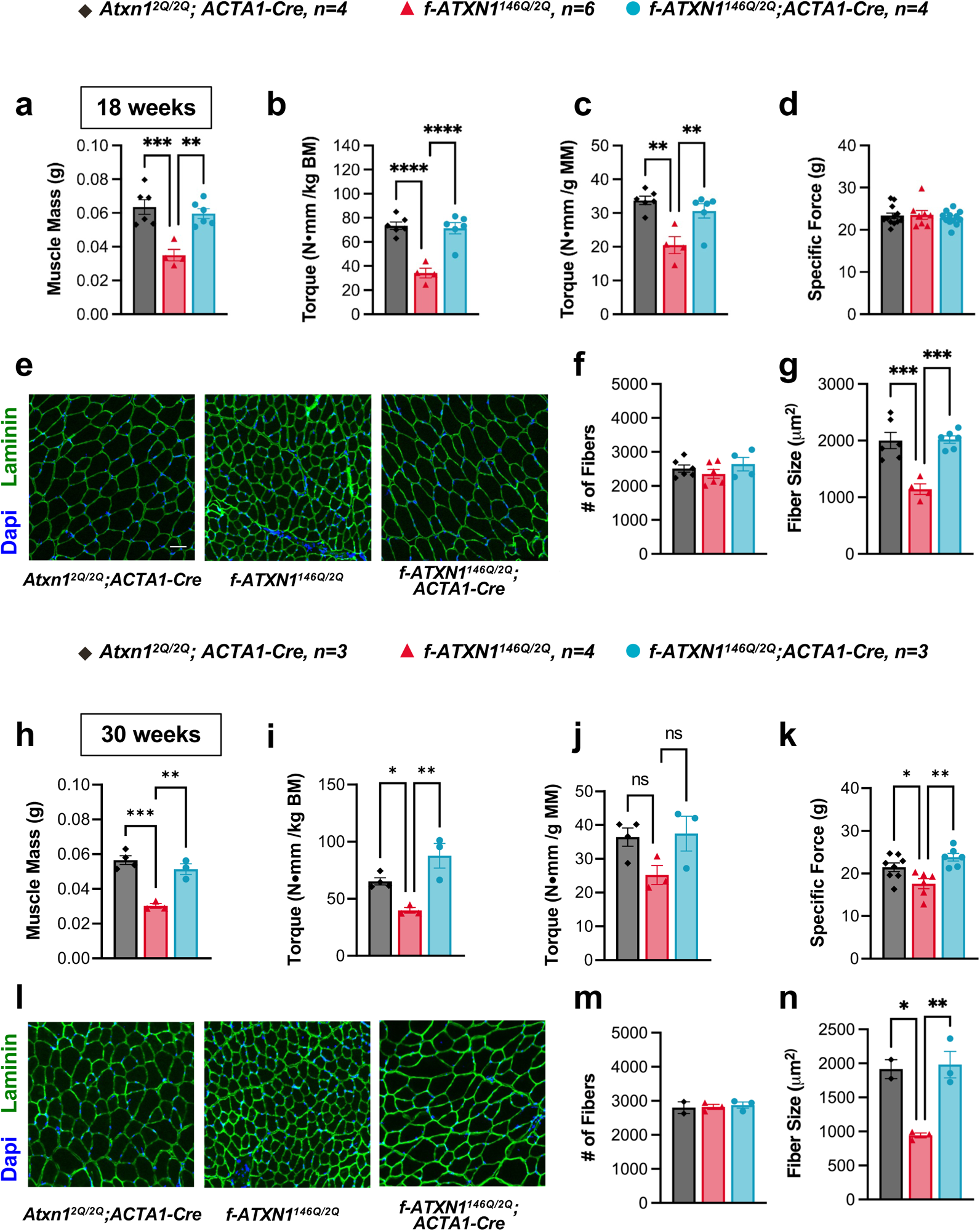
Skeletal muscle pathology in *f-ATXN1^146Q/2Q^* is corrected with deletion of the *ATXN1^146Q^* allele. **a, h,** Dorsiflexor muscle mass comprised of the sum of tibialis anterior (TA) and extensor digitorum longus (EDL) muscle masses for 18 and 30-week-old *Atxn1^2Q/2Q^; ACTA1-Cre*, *f-ATXN1^146Q/2Q^*, and *f-ATXN1^146Q/2Q^; Acta1-Cre* mice. **b-c, i-j,** Peak anterior crural isometric torque at 18 and 30 weeks normalized to body mass and dorsiflexor mass. **d, k,** Specific isometric force from *ex vivo* preparations of EDL muscles at 18 and 30 weeks. **e, l,** Representative cross-sectional images from TA muscles showing nuclear (blue; DAPI) and laminin (green) staining at 18 and 30 weeks. **f, m,** Quantification of total number of fibers and **g,n,** average fiber size for TA muscle cross-sections at 18 and 30 week. One-way ANOVAs with Tukey’s post hoc test were performed for all except **e** and **l**. Significance of results is denoted as * (p<0.05), ** (p<0.01), *** (p<0.001).

**Figure 6.**
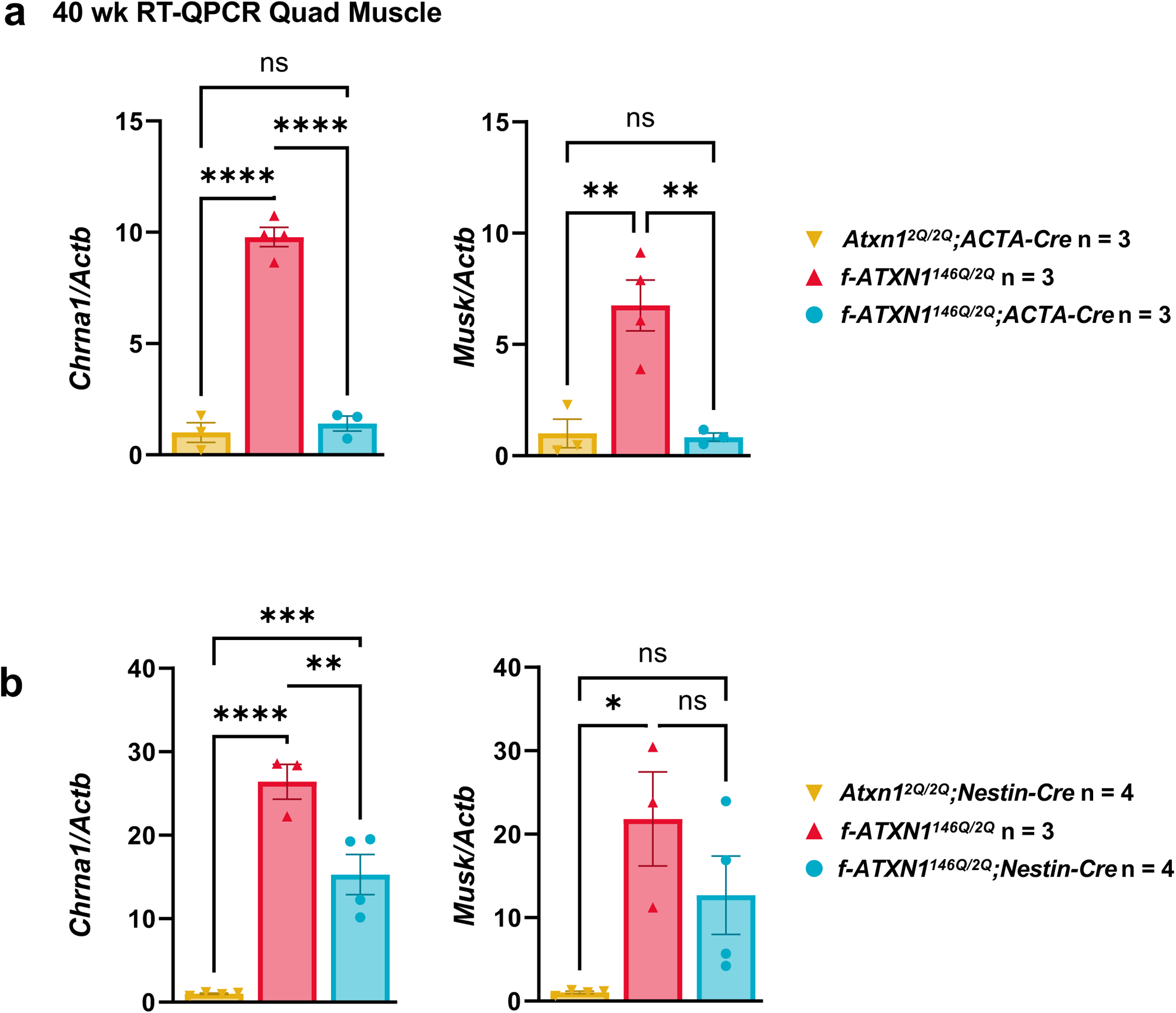
NMJ mRNAs qPCR. **a,** Relative expression of *Chrna1* and *Musk* in 40 wk quadricep RNA from *Atxn1^2Q/2Q^; ACTA1-Cre*; *f-ATXN1^146Q/2Q^*, and *f-ATXN1^146Q/2Q^*mice. **b,** Relative expression of *Chrna1* and *Musk* in 40 wk quadricep RNA from *Atxn1^2Q/2Q^; Nestin-Cre*; *f-ATXN1^146Q/2Q^*, and *f-ATXN1^146Q/2Q^*mice. One-way ANOVAs with Tukey’s post hoc test were performed for all. Significance of results is denoted as * (p<0.05), ** (p<0.01), *** (p<0.001), **** (p<0.0001).

Histological analysis of muscle cross sections at 18 and 30 weeks revealed that total fiber number of the TA was not different between genotypes, but the *f-ATXN1^146Q/2Q^* TA muscles had significantly smaller fiber cross-sectional area compared to *Atxn1^2Q/2Q^;ACTA1-Cre* TA muscles (Figures 5e-5g and 5l-5n, respectively). Muscle fiber sizes were completely rescued in *f-ATXN1^146Q/2Q^*; *ACTA 1-Cre* mice (Figures 5g and 5n). These data support the presence of a *f-ATXN1^146Q/2Q^* specific skeletal muscle pathology that is characterized by muscle weakness and atrophy.

Proper nuclear localization of mutant ATXN1 is critical for many disease-like phenotypes including motor dysfunction, cognitive deficits, and premature lethality^23^. Like in *f-ATXN1^146Q/2Q^* mice, smaller dorsiflexor muscle mass (Figure 7a), and smaller fiber size (Figure 7g) indicating the presence of a skeletal muscle myopathy are seen in *Atxn1^175Q/2Q^* mice as early as 12 weeks-of-age. As in *f-ATXN1^146Q/2Q^* mice, *Atxn1^175Q/2Q^* mice had the same number of muscle fibers as *Atxn1^2Q/2Q^* mice (Figure 7f). Interestingly, this phenotype was partially corrected in *Atxn1^175Q-K772T/2Q^* mice. The *Atxn1^175Q-K772T/2Q^* muscles demonstrate partially recovered muscle mass (Figure 7a), and muscle fiber size (Figures 7e and 7g). Thus, as with the other SCA1-like disease phenotypes, we conclude that nuclear localization of expanded ATXN1 is important for the muscle-specific phenotypes caused by expanded ATXN1.

**Figure 7.**
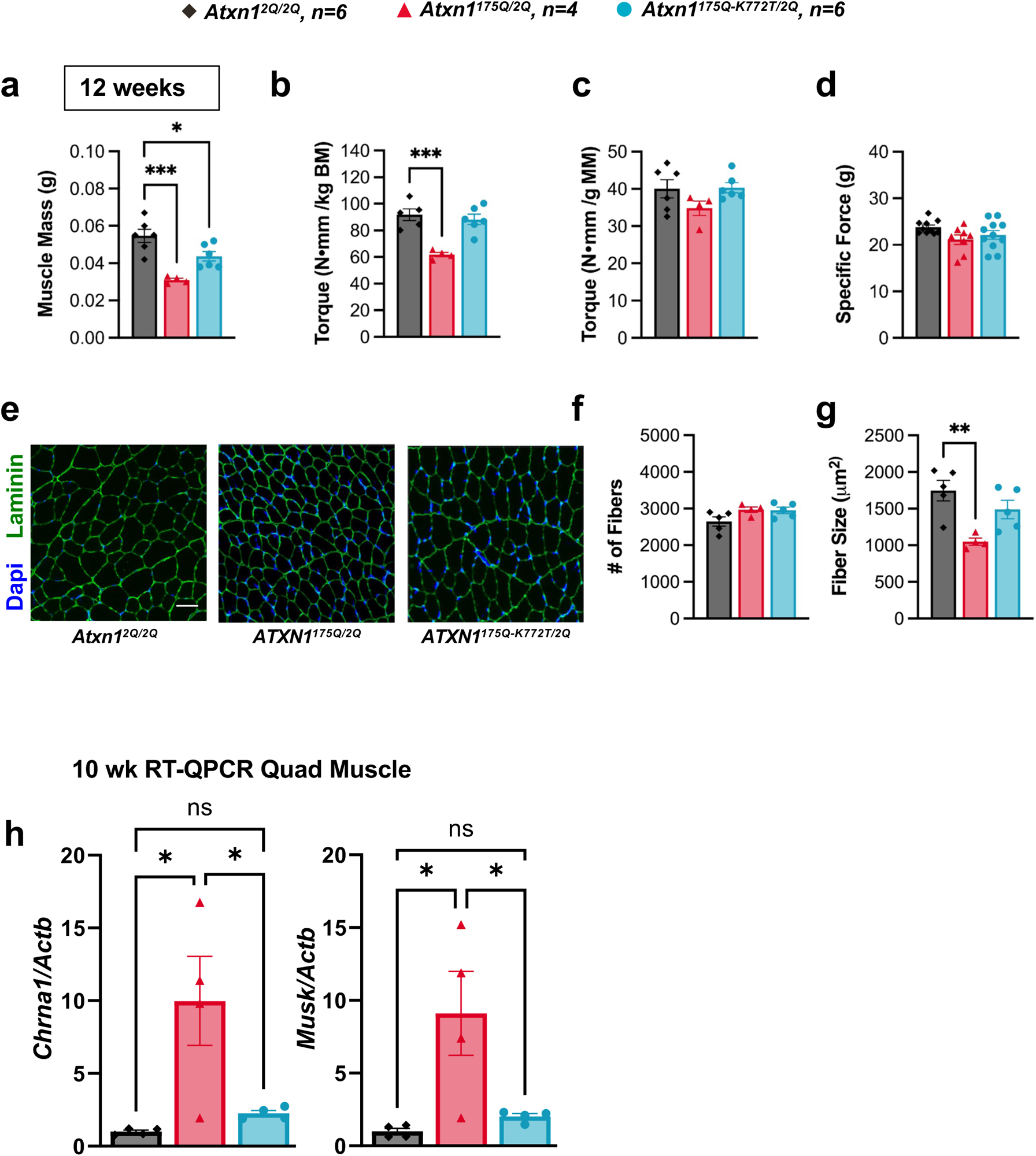
Nuclear localization of expanded ATXN1 is important for muscle-specific phenotypes caused by expanded ATXN1. **a,** Dorsiflexor muscle mass comprised of the sum of tibialis anterior (TA) and extensor digitorum longus (EDL) muscle masses for 12-week-old *Atxn1^2Q/2Q^*, *Atxn1^175Q/2Q^* and *Atxn1^175Q-K772T/2Q^* mice. **b,** Peak anterior crural isometric torque normalized to body mass and **c,** dorsiflexor mass. **d,** Specific isometric force from *ex vivo* preparations of EDL muscles. **e,** Representative cross-sectional images from TA muscles showing nuclear (blue; DAPI) and laminin (green) staining. **f,** Quantification of total number of fibers and **g,** average fiber size from TA muscle cross-sections. **h,** Relative expression of *Chrna1* and *Musk* in 40 wk quadricep RNA from *Atxn1^2Q/2Q^*, *Atxn1^175Q/2Q^* and *Atxn1^175Q-K772T/2Q^* mice. One-way ANOVAs with Tukey’s post hoc test were performed for all except **e**. Significance of results is denoted as * (p<0.05), ** (p<0.01), and *** (p<0.001).

Figures S7 depicts the impact of deleting the *f-ATXN1^146Q^* allele from muscle has on additional SCA1-like phenotypes. *F-ATXN1^146Q/2Q^*; *ACTA1-Cre* mice showed no improvement in open field, rotarod and hind limb clasping phenotypes (Figures S76a-S7c). In contrast, survival and wasting were significantly improved in *f-ATXN1^146Q/2Q^*; *ACTA1-Cre* mice (Figures S7d & S7e) supporting the concept that mutant ATXN1 induced muscle pathology contributes to these SCA1-like phenotypes.

## Discussion

Neurodegenerative diseases are often characterized by prominent pathology in specific brain regions/neuronal populations, yet many of these disorders impact other neural structures, circuits, and cell populations that lack obvious pathology. In SCA1, cerebellar Purkinje cell degeneration is a frequent and prominent pathological feature. However, ATXN1 is widely expressed and SCA1 patients present with symptoms linked to multiple different brain regions^7^. A question facing the understanding of SCA1 pathogenesis, as well as many other neurodegenerative diseases, is the extent to which vulnerability in specific cell populations contribute to each disease-associated phenotype. In this study, we addressed this question in SCA1 using a conditional mouse model with broad spatial mutant gene expression that matches that of humans. We found that deletion of the *f-ATXN1^146Q^* allele from CNS neuronal and glial cells rescued SCA1-like neurological phenotypes seen in the *f-ATXN1^146Q/2Q^* mice. Analysis of a key disease phenotype, deficit in motor performance on the rotarod, showed that this phenotype stems from effects of *f-ATXN1^146Q^* in at least one brain region other than cerebellar Purkinje cells, i.e. MSNs in the striatum. Additionally, direct pathological effects of *f-ATXN1^146Q^* on muscle were found.

Like the *Atxn1^154Q/2Q^* knockin mice^8^, *f-ATXN1^146Q/2Q^* mice developed neurological phenotypes similar to those seen in SCA1 patients. Motor incoordination on the rotarod was seen as early as 7 weeks of age in *f-ATXN1^146Q/2Q^* mice and cognitive deficits as assessed by the Barnes maze test later at 24 weeks of age. Confirming the neural basis of these SCA1-like phenotypes, substantial rescue of rotarod performance (Figure 3b), performance in the open field test (Figure 2a), and on the Barnes maze (Figure 2b) was observed in *f-ATXN1^146Q/2Q^* mice crossed to *Nestin-Cre* mice. The *Nestin-Cre* line was designed to direct Cre recombinase expression to neuronal and glial precursors providing recombination exclusively in the CNS^10^. Analysis of the tissue pattern of *f-ATXN1^146Q^* deletion in *f-ATXN1^146Q/2Q^; Nestin-Cre* mice confirmed the CNS specificity of *Nestin-Cre* recombination (Figures S3a,S3b & S3d). As an initial effort to further dissect the anatomical basis of a key SCA1-like phenotype manifested by the *f-ATXN1^146Q/2Q^* mice, we selected motor incoordination as measured by rotarod performance. Interestingly, while injection of an *Atxn1* targeting RNA into the cerebellum was able to improve rotarod performance at 10 weeks of age in *Atxn1^154Q^* knockin mice^12^, deletion of *f-ATXN1^146Q^* from the striatum by crossing *f-ATXN1^146Q/2Q^* mice to *Rgs9-Cre* mice rescued rotarod performance late in disease progression at 31 weeks of age. Correspondingly at diagnosis, magnetic resonance imaging (MRI) of SCA1 patients shows that loss of cerebellar volume is essentially complete. As SCA1 patients age, MRI analyses shows a progressive loss in striatal volume^14,15^. Moreover, a recent study found striatal volume to be a predictor of motor decline with increasing patient age after onset of ataxia^15^. Our findings indicate that a relative time course of disease in cerebellum and striatum in *f-ATXN1^146Q/2Q^* mice parallel the MRI findings in SCA1 patients. It is intriguing that there is somatic instability of expanded *ATXN1* in the striatum of both SCA1 knockin mice and SCA1 patients^18,19,24^ (Figures 3d and 3e). Here, we show that in *f-ATXN1^146Q/2Q^* mice, ATXN1 CAG expansion occurs in MSNs (Figures 3d and 3e). We speculate that striatal cells are less sensitive to expanded ATXN1 than Purkinje cells, requiring the somatic expansion with age for initiation of striatal pathogenesis. Somatic expansion in other brain regions/cell-types (Figure 3d) may also contribute to disease pathogenesis. Regardless, it is clear that regions/cell populations in addition to cerebellum/ Purkinje cells, and specifically the striatum/msn’s, contribute to the rotarod deficit in *f-ATXN1^146Q/2Q^* mice.

As disease progresses in SCA1 patients, muscle wasting associated with severe weight loss is often seen^25-28^. In one study, it was noted that in some cases wasting seemed not to be due to dysphagia or altered by diet^28^. Signs of motor neuron pathology were suggested as being linked to muscle wasting^29^. In this study, we provide evidence that expanded ATXN1 has a direct pathological effect on skeletal muscle. At 18 weeks we observed that *in vivo* torque deficits were not corrected even after normalizing muscle strength by muscle size. This suggests a muscle force-generating pathology that could be stemming from a variety of muscle contraction-related elements ranging from motor neuron activation to myosin-actin cross bridge cycling. Interestingly, *ex vivo* force at 30 weeks was significantly lower in *f-ATXN1^146Q/2Q^* mice than *Atxn1^2Q/2Q;^ACTA1-Cre* mice, which suggests that the muscle pathology could progress to more specific disruptions in intrinsic contractile function. Notably, every muscle deficit was restored to that seen in WT *Atxn1^2Q/2Q^* mice in *f-ATXN1^146Q/2Q^* mice in which *ACTA1-Cre* induced deletion of the *ATXN1^146Q^* allele. Overall, the data point to the presence of a neuromuscular pathology perhaps involving dysfunction at the NMJ. That NMJ function is impaired in *f-ATXN1^146Q^*mice is supported by the finding that expression of *MuSK* and *Chrna1* RNA is altered in muscle of *f-ATXN1^146Q^* mice. Further investigation is necessary to explore the contractile-specific effects of the muscle-specific ATXN1-[146Q] mutation.

Notably, deletion of *f-ATXN1^146Q^* from muscle substantially reduced kyphosis that in mice, is associated with paraspinal skeletal muscle atrophy^20^ as well as restored normal dorsiflexor strength, muscle mass and fiber size. As seen for other SCA1-like phenotypes, motor dysfunction, cognitive deficits, and premature lethality^23^, proper nuclear localization of expanded ATXN1 was shown to be critical for muscle pathogenesis. These results show that *f-ATXN1^146Q/2Q^* along with *f-ATXN1^146Q/2Q^*; *Acta1-Cre* will provide an excellent experimental platform for elucidating the molecular aspects of ATXN1[146Q]-induced muscle pathogenesis.

In conclusion, our study demonstrates the utility of the *f-ATXN1^146Q/2Q^* conditional mouse model in linking pathogenesis in specific anatomical regions/cell populations with SCA1-phenotypes. The results reveal that neural and peripheral pathological effects of expanded ATXN1 contribute to disease that have important implications for design and administration of optimal SCA1 therapeutics. Notably, we show that the rotarod motor performance deficit is more complex in origin than just pathology of cerebellar Purkinje cells and that another critical region is the striatum as *f-ATXN1^146Q/2Q^* mice age. Lastly, data presented provide strong evidence of muscle specific pathology in *f-ATXN1^146Q/2Q^* mice. Thus, an ideal SCA1 therapeutic should target pathogenic effects of mutant ATXN1 throughout the brain including cerebellar Purkinje cells and the striatum as well as other brain regions. In addition, an optimal SCA1 treatment needs to affect mutant ATXN1 toxicity in peripheral muscle.

## ACKNOWLEDGEMENTS

This study was supported by NIH/NINDS grants R01NS022920, R35NS127248 and NS049206. The authors thank the Yun You and the Mouse Genetics Laboratory at the University of Minnesota and the MGH Mission Driven Service Core for DNA Fragment Analysis.

## AUTHOR CONTRIBUTIONS

L.D. and H.T.O. conceived the study and wrote the paper. L.D., W.M.S., J.M.E., M.C. and H.T.O. designed experiments and interpreted the data. K.B. and M.D.K designed the strategy and generated the *f-ATXN1^146Q/2Q^*mouse model. Y.Y. performed embryo injections and implantations. O.R. and S.S. bred mice and managed the colony. L.D., O.R., and S.S. conducted the survival study and animal weight measurements. O.R., E.L. and T.N.-M. performed behavioral assays. H.K. performed qRT-PCR, western analyses and genotyping. P.Y., A.S., and C.S., performed dissection, and recombination assay. U.K.G. and B.O’C. assessed degree of Cre-mediated recombination. B.O’C performed immunofluorescence and imaging. J.S.M. developed and implemented kyphosis image analyses. W.M.S. performed muscle histological and functional analyses. H.P.H. generated the *Atxn1^175QK772T/2Q^* mouse model and performed statistical analyses. H.Y.Z. designed experiments and edited the manuscript. V.C.W, H.T.O and L.D. designed the repeat instability experiment. Z.N.B performed the repeat instability experiment and analyzed the data. V.C.W and Z.N.B interpreted the data and V.C.W contributed to the writing of the manuscript. All authors reviewed the manuscript and provided input.

---

**Figure S1.**
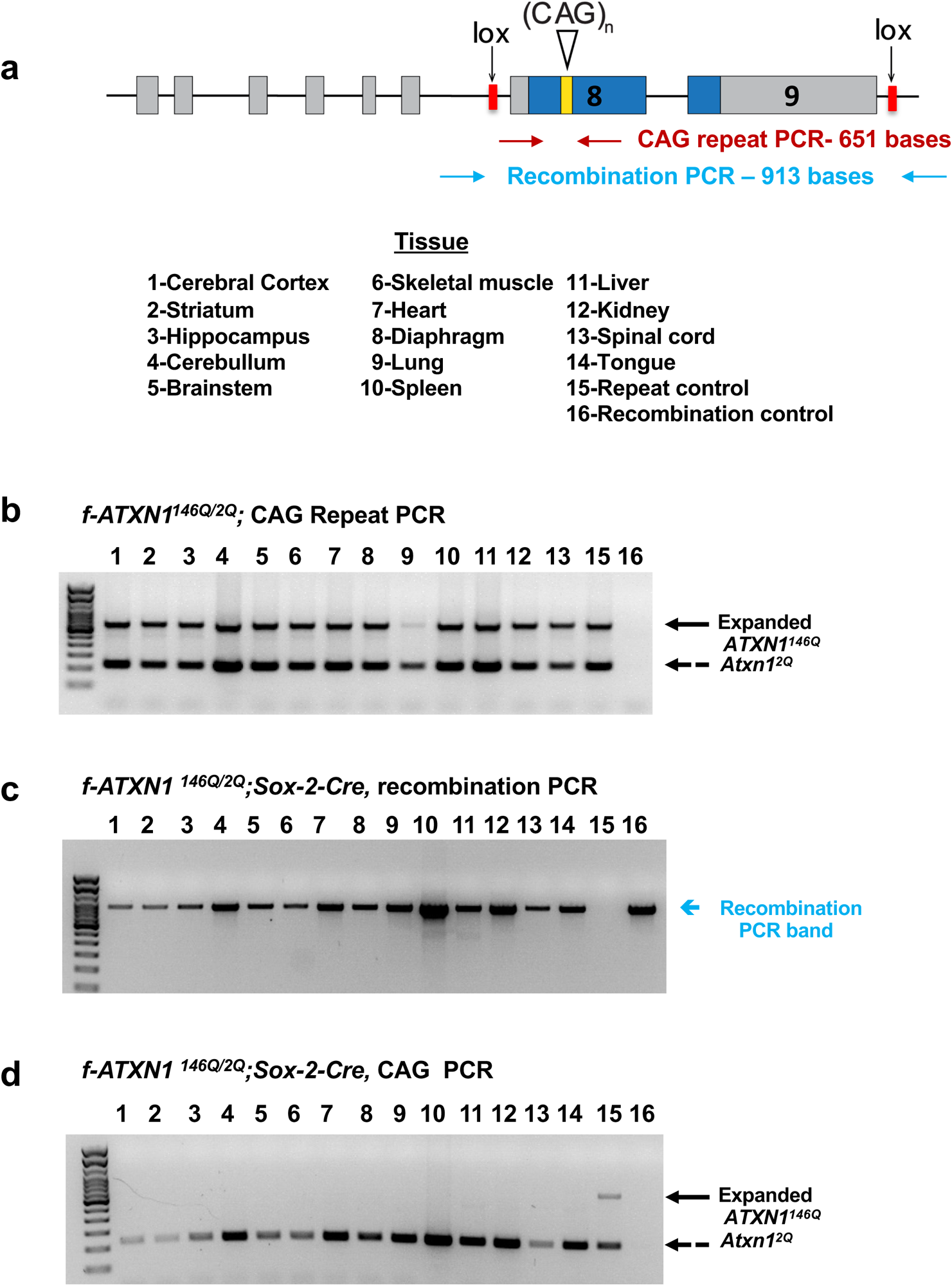
*F-ATXN1^146Q/2Q^* diagram and PCR. **a,** Schematic of *f-ATXN1^146Q^* allele into the mouse endogenous locus. **b,** Repeat PCR in tissue DNA from the *f-ATXN1^146Q/2Q^***. c,** Repeat PCR in tissue DNA from *f-ATXN1^146Q/2Q^*; *Sox-Cre***. d,** Recombination PCR in tissue DNA from *f-ATXN1^146Q^*; *Sox-Cre*.

**Figure S2.**
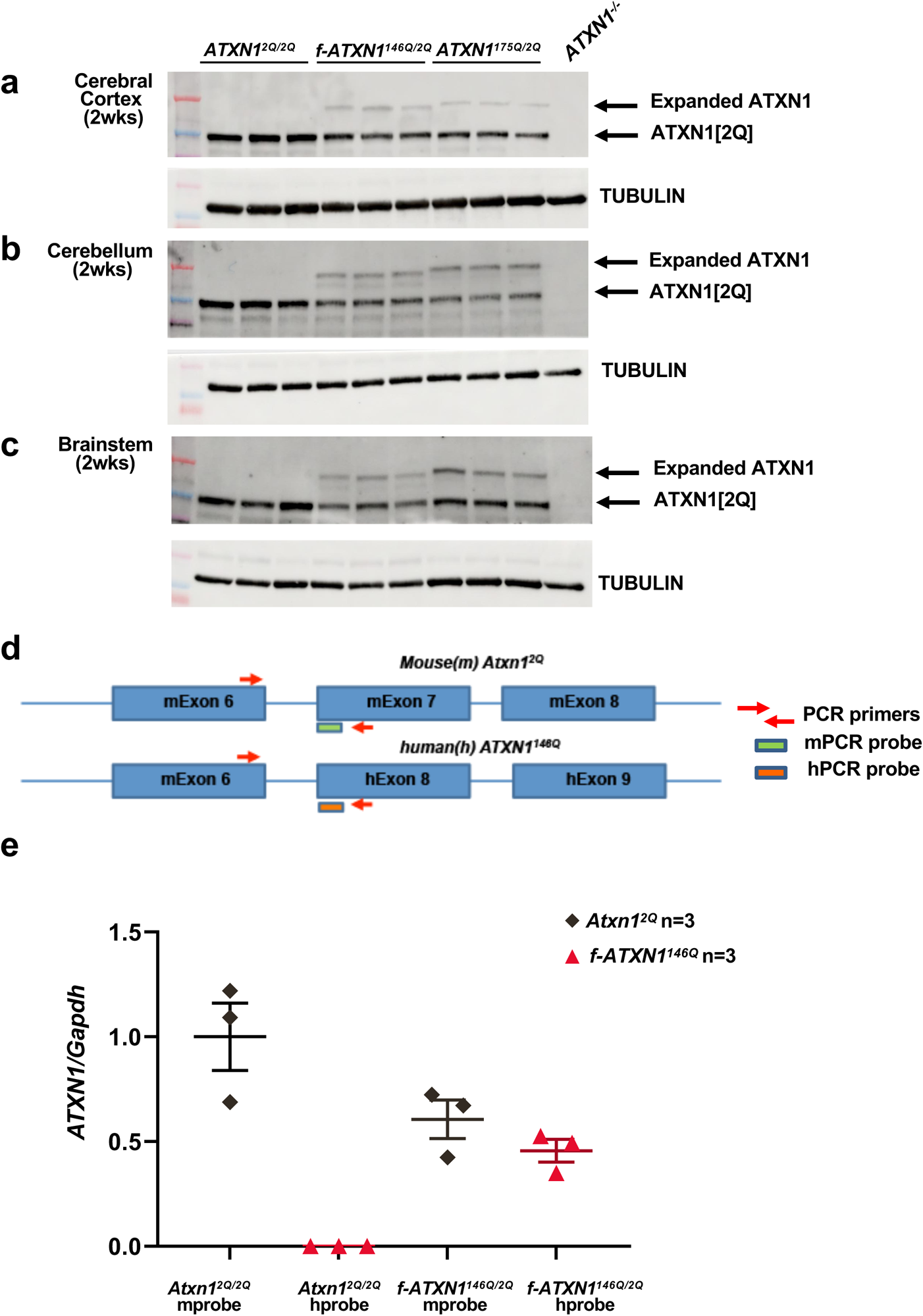
*F-ATXN1^146Q/2Q^* mice express equal protein and RNA as endogenous gene. **a-c,** Western blot of expanded ATXN1 in *f-ATXN1^146Q/2Q^* and *Atxn1^175Q/2Q^* mice from the cortex, cerebellum, and brainstem in 2-week-old mice. **d,** Schematic of the RT-qPCR reaction designed to quantify either *hATXN1* or *mAtxn1* RNA expression using identical primer sequences and unique probe sequences. **e,** Relative expression of *hATXN1* and *mAtxn1* in *Atxn1^2Q/2Q^* mice and *f-ATXN1^146Q/2Q^* mice.

**Figure S3.**
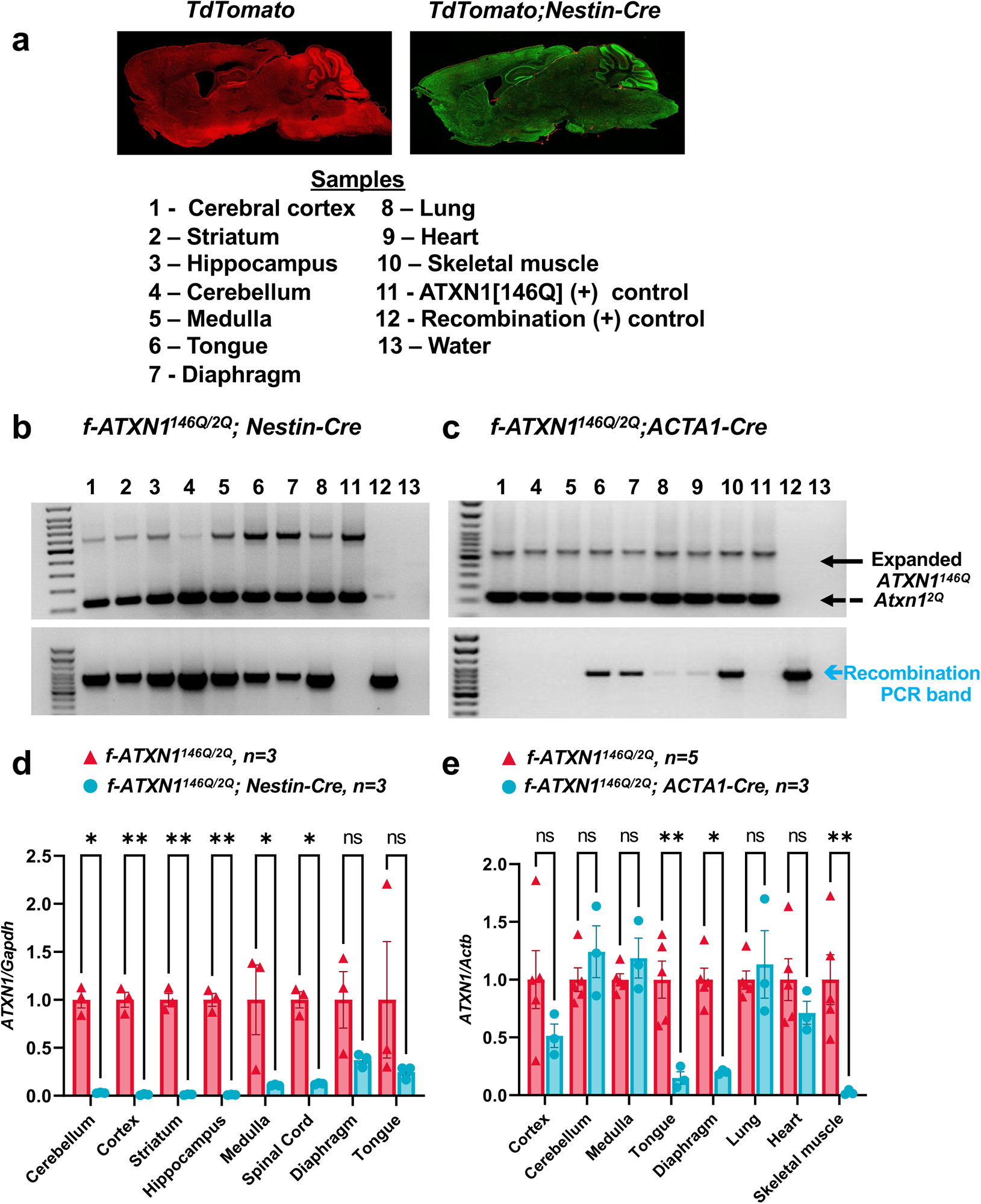
Human *ATXN1* expression in *f-ATXN1^146Q^;Nestin-Cre* and f-ATXN1^146Q^;ACTA1-Cre *mice*. **a,** Representative images showing Cre-recombination from a *TdTomato;Nestin-Cre* reporter mouse (green). **b,** Repeat and recombination PCR in tissue DNA from the *f-ATXN1^146Q^; Nestin-Cre* and **c,** *f-ATXN1^146Q^; ACTA1-Cre* mouse**. d,** RT-qPCR of ATXN1 knockdown in tissue from *f-ATXN1^146Q^; Nestin-Cre* and **e,** *f-ATXN1^146Q^; ACTA1-Cre* mice. Two-way ANOVA with Šídák post hoc test. Significance of results is denoted as * (p<0.05) and ** (p<0.01).

**Figure S4.**
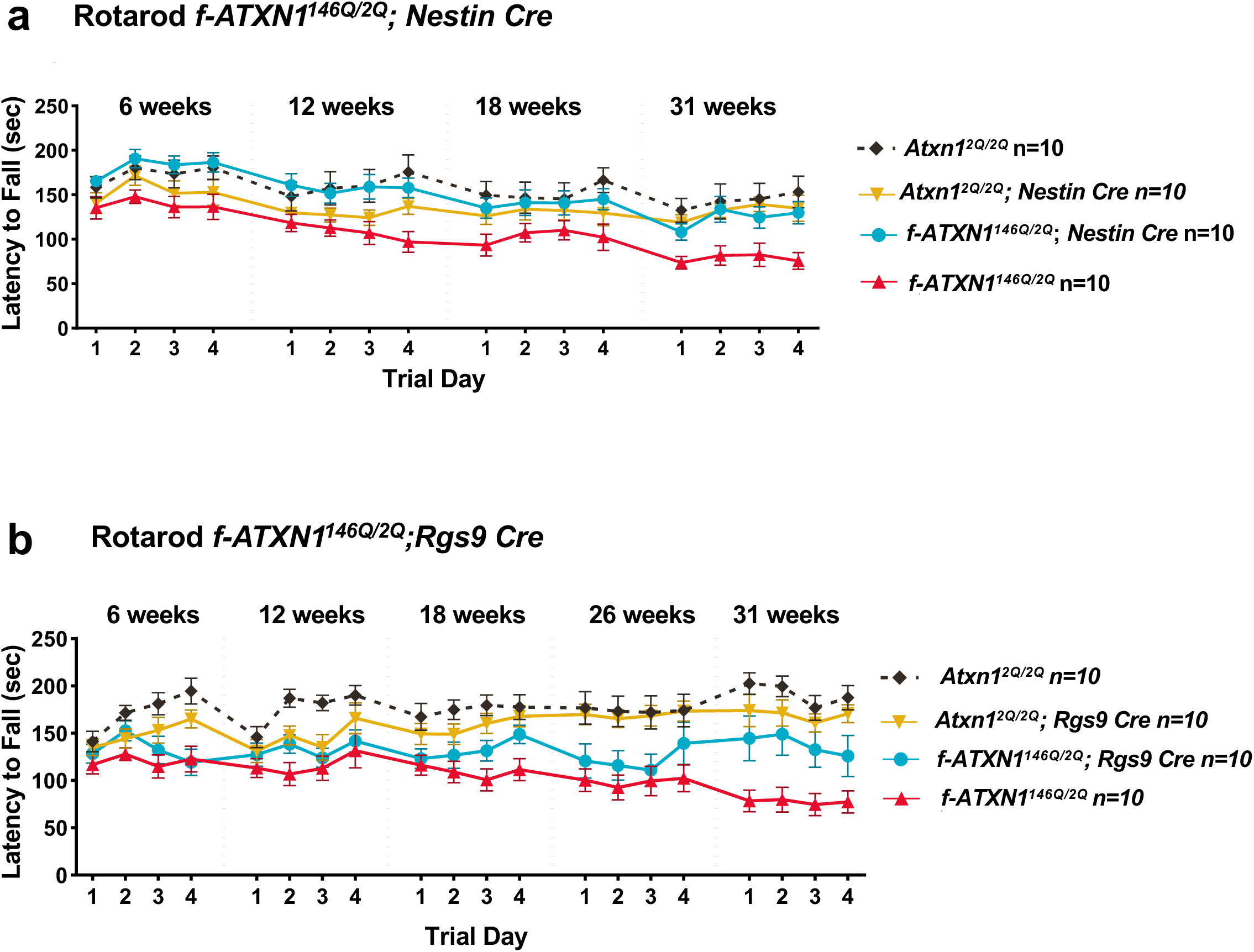
CNS regional contribution to progressive motor performance deficit in *f-Atxn1^146Q/2Q^* mice. **a** & **b,** Rotarod assessment of all 4 genotypes.

**Figure S5.**
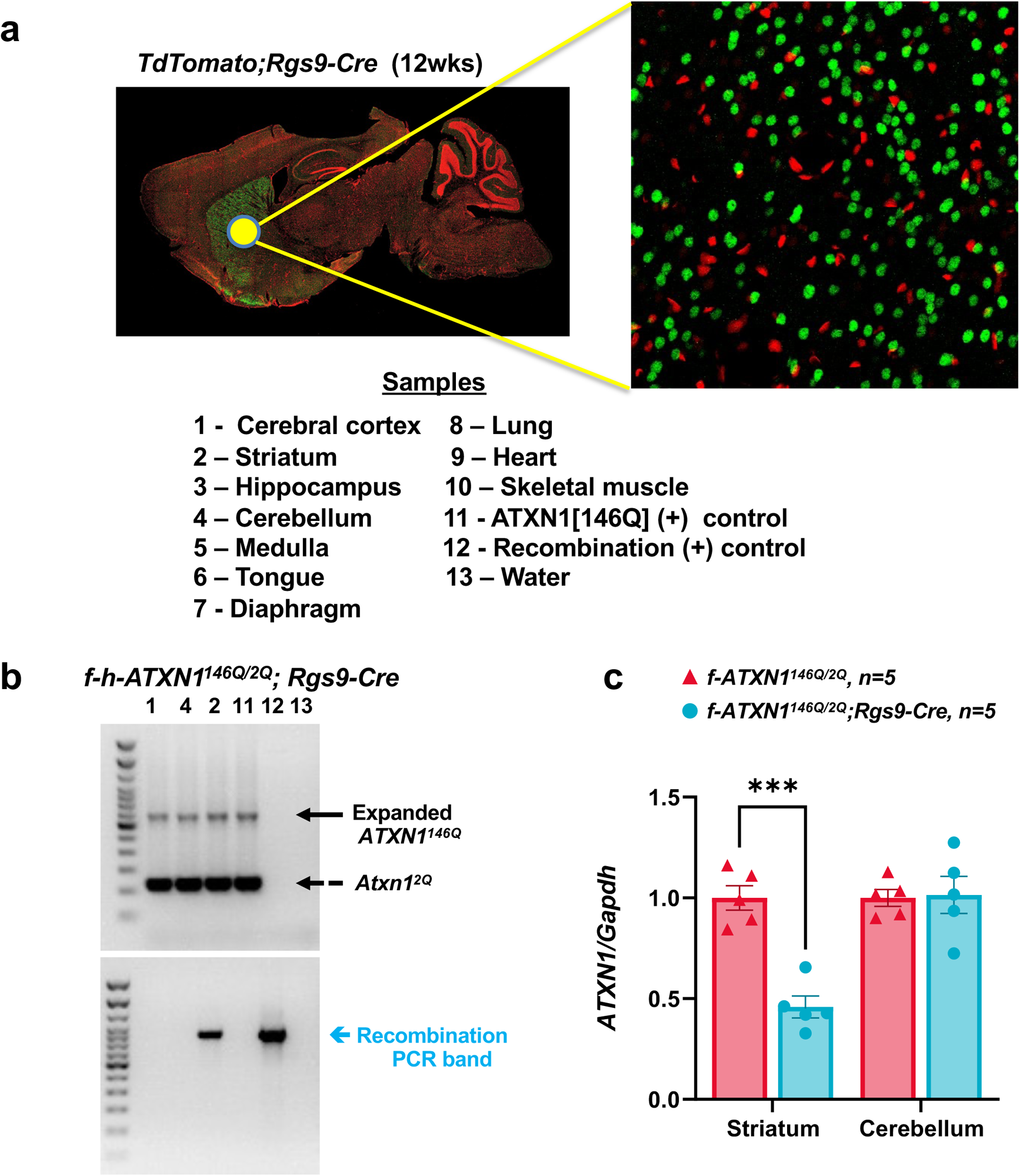
Human *ATXN1* expression in *f-ATXN1^146Q^; Rgs9-Cre* mice. **a,** Representative images showing Cre-recombination in the striatum (green) from a *TdTomato; Rgs9-Cre* reporter mouse. **b,** Repeat and recombination PCR in tissue DNA from the *f-ATXN1^146Q^; Rgs9-Cre*. **c,** RT-qPCR of ATXN1 knockdown in tissue from *f-ATXN1^146Q^; Rgs9-Cre*. Two-way ANOVA with Šídák post hoc test. Significance of results is denoted as *(p<0.05), ** (p<0.01), and *** (p<0.001).

**Figure S6.**
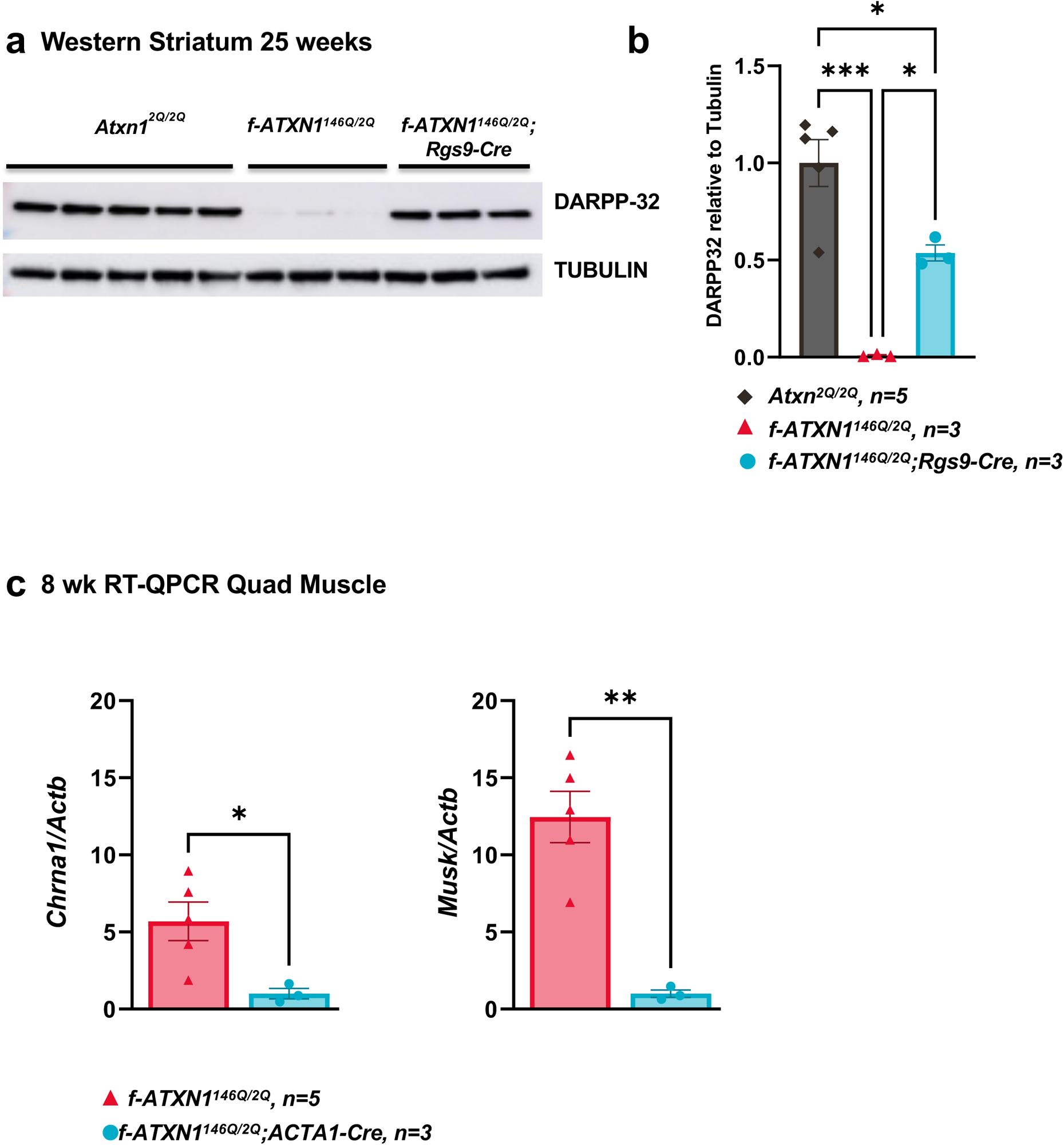
**a**, Western blot for quantification of DARPP-32 protein from striatum of *Atxn1^2Q/2Q^, f-ATXN1^146Q/2Q^, f-ATXN1^146Q/2Q^;Rgs9-Cre* mice. **b,** Quantification of DARPP32 protein relative to TUBULIN. One-way ANOVA with Tukey post hoc test **c,** Relative expression of *Chrna1* and *Musk* in 8 wk quadricep RNA from *Atxn1^146Q/2Q^*and *f-ATXN1^146Q^; ACTA1-Cre.* One-way ANOVAs with Tukey’s post hoc test were performed for all. Significance of results is denoted as * (p<0.05), ** (p<0.01), *** (p<0.001).

**Figure S7.**
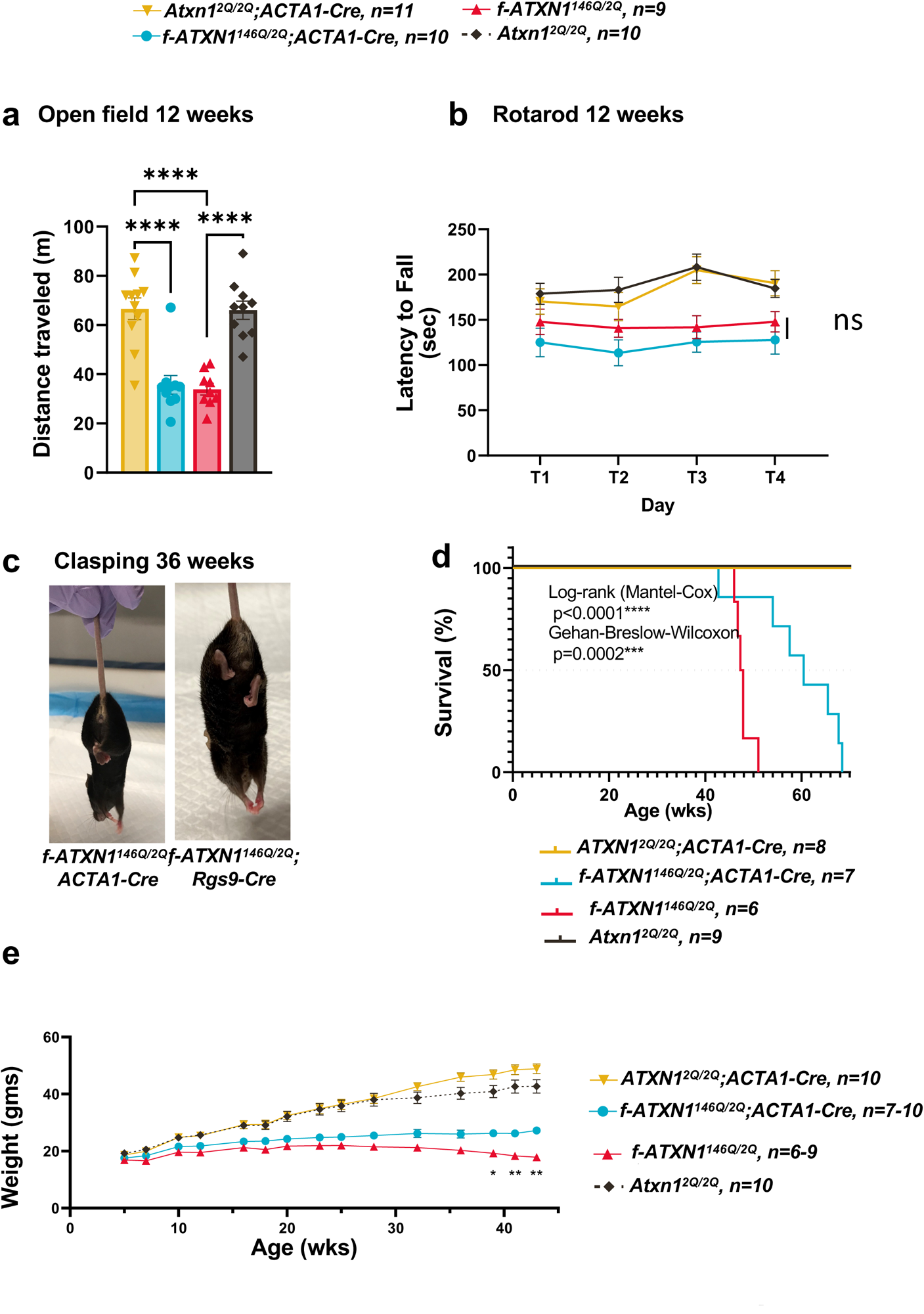
SCA1-like phenotypes in *f-ATXN1^146Q/2Q^*; *ACTA1-Cre* mice. **a,** Distance traveled in open field at 12 weeks of age. One-way ANOVA with Tukey post hoc test. **b,** Rotarod assessment of all 4 genotypes at 12 weeks. No significant difference between *f-ATXN1^146Q/2Q^; ACTA1-Cre* and *f-ATXN1^146Q/2Q^*. Two-way ANOVA with Tukey post hoc test. **c,** Clasping phenotype in *f-ATXN1^146Q^; ACTA1-Cre,* and *f-ATXN1^146Q^;Rgs9-Cre*. **D,** Mouse survival plotted as Kaplan-Meyer curves with median lifespan labeled for each genotype. Log-rank Mantel-Cox and Gehan-Breslow-Wilcoxon test. **e**, Body weight measurements between 5 and 45 weeks of age. Two-way ANOVA Tukey post hoc test. Significance of results is denoted as * (p<0.05), ** (p<0.01), *** (p<0.001) and ****(p<0.0001).

## Methods

### Mice

The University of Minnesota Institutional Animal Care and Use Committee approved all animal use protocols. All mice were housed and managed by Research Animal Resources under specific pathogen-free conditions in an Association for Assessment and Accreditation of Laboratory Animal Care International approved facility. The mice had unrestricted access to food and water except during behavioral testing. In all experiments, equal numbers of male and female mice were used. All mice were age matched within experiments and littermate controls were used when possible. All mice were maintained on a C57BL/6 genetic background. *Atxn1^2Q/2Q^* (C57Bl6/J) mice, *ACTA1-Cre* (B6.Cg-Tg(ACTA1-cre)79Jme/J RRID:IMSR_JAX:006149) mice, and *Nestin-Cre* (B6.Cg-Tg(Nes-Cre)1Kln/J RRID:IMSR_JAX:003771), *TdTomato* (B6N.129S6-Gt(ROSA)26Sor^tm1(CAG-tdTomato*,-EGFP*)Ees^/J RRID:IMSR_JAX:023537) mice were obtained from The Jackson Laboratory. The *Rgs9-Cre* mice were a generous gift from Dr. X. William Yang, University of California Los Angeles.

### Generation of *f-ATXN1^146Q/2Q^* mice

Embryonic stem cell culture: C57BL/6N-PRX-B6N mouse embryonic stem (mES) cells were purchased from Jackson Laboratories (donating investigator Robin Wesselschmidt, Primogenix, Inc). Cells were maintained at 37°C at 7.5% CO_2_ in IMDM (Iscoves DMEM) media containing 20% ESC qualified FBS, 1X NEAA, 2mM L-glutamine, 1X Penicillin/Streptomycin, 0.2mM Beta-Mercaptoethanol, 1000 U/ml ESGRO-mouse LIF.

Cells were grown on gelatin coated dishes containing irradiated mouse embryonic fibroblasts (iMEF). mES cell transfections, selection, analysis and expansion: Genomic recombination was enhanced by Cas9-CRISPR cleavage at a site 5’ of the mouse *Atxn1* exon 7 (5’-TAAGCGGCTGTCTTGACCAC) and a site 3’ of the *Atxn1* polyA site (5’-TAAGCTGTGGTTGCTTGAGC) 19.6kb from the first cleavage site. These sgRNA targets were cloned into pSpCas9(BB)-2A-Puro (AddGene plasmid ID 48139) as described.^30^ mES cells were trypsinized with 0.05% trypsin, plated onto gelatin coated (no iMEF) 6 well dishes from a 90% ESC confluent p60 dish at a 1:6 dilution, and transfected with the pair of sgRNA/Cas9 plasmids and a repair template that inserted between the two cleavage sites a puromycin cassette flanked by FRT sites and an adjacent, promoterless neomycin gene that ended in a Lox site (Fig. 1aii). The next day, the ESC were transferred to 10cm, gelatin coated iMEF-puroR plates. Cells were allowed to recover for 24 hours. 1.4ug/mL puromycin was added on day 2 and 3, with no puromycin on day 4. 1.5ug/mL puromycin was added again on days 5-7. On day 8, selected clones were picked onto 96 well gelatin iMEF plates. When cells were 80-90%confluent, ES cells were frozen in duplicate 96 well format (80% compete media, 10% additional ES FBS, 10% DMSO) and a third onto 48 well gelatin-only plate, in which cells were allowed to expand and harvested for DNA analysis of each clone by PCR with the following primers: i6Atxn1 Junction (F1: 5’-ACACGTGGCTGCAATTTGTC; R1: 5’-GTAACGCGCTTGCTGCTTG) and 3’ of Atxn1 Junction (F1: 5’-AGCGTATCCACATAGCGT; R1: 5’-CTTGCCCATTGCATACCAGG). The puromycin cassette in clone 40 from this first transfection was replaced by human genomic DNA syntenic to the deleted mouse genomic sequences by Flp recombinase using the approach as described^31^. This 31kb of human sequence (BAC RP11-413J6) extends from sequences 5’ of human ATXN1 exon 8 (5’-TAATGTTACACCAGGCTAAA) to a sequence 3’ of the human ATXN1 polyA site (5’-AGGTGGAATCCCCTGCACCC), and is flanked by FRT sites, a Lox sequence just 3’ of the 5’ FRT site, contains a 146Q expanded CAG repeated in exon 8, and at the 3’ end has a promoter-FRT cassette that drives expression of the neomycin resistance gene upon recombination into the modified mES cell. After transfection, mES cells were transferred to 10cm, gelatin coated iMEF-Neo plates. Cells were allowed to recover for 48 hrs and were then selected with 125ug/mL of G418 (50mg/mL Gibco) for 7 days. Selected clones (13 total) were picked and expanded. The clones were frozen with 80% complete media, 10% additional ES-FBS, and 10% DMSO. PCR analyses were performed with the following primers: i6Atxn1 Junction (F1: 5’-ACACGTGGCTGCAATTTGTC; R2: 5’-GGATGGCTCTGATTTTAGTCTG) and NeoR Junction (F1: 5’-ACGAGCCTTCATAGCATCCG; R1: 5’-GGATGGCTCTGATTTTAGTCTG). Clone 2 and C10 were correct for all assays and sequence verified and further expanded. Chimeric mice were generated by injection of C2 and C10 ESC into blastocysts and backcrossed with C57BL/6J.

### Genotyping

PCR was performed with the following primers (Integrated DNA Technologies) to determine which animals have an expanded *f-ATXN1^146Q/2Q^* allele: ATXN1-146Q repeat Forward (5’-CAACATGGGCAGTCTGAG) and ATXN1-146Q repeat Reverse (5’-GTGTGTGGGATCATCGTCTG) To assess recombination by Cre the following primer set was used: Recombination Forward (5’-GGGAATGGTACCAACCTT TCTG) and Recombination Reverse (5’-GTAGAACCCCAGACCCTCGT). Cre F oIMR 1084 (5’-GCGGTCTGGCAGTAAAAACTATC) and Cre R oIMR 1085 (5’-GTGAAACAGCATTGCTGTCACTT) were used to genotype all Cre lines.

### Analyses of CAG instability

Genomic DNA was extracted from dissected striatum, medulla, cortex, hippocampus and cerebellum using the Qiagen DNeasy Blood & Tissue kit according to the manufacturer’s protocol. Tail DNA was prepared using Wizard SV Genomice DNA kit (Promega, A2360). The *ATXN1* CAG repeat was PCR-amplified using 1μM each forward (6-FAM-5^′^-CAGAGTGGAATAGGCCTCCA) and reverse (5’-TGGACGTACTGGTTCTGCTG) human *ATXN1*-specific primers, 10 µl 2X Promega GoTaq Colorless Master Mix and 4 µl betaine in a total reaction volume of 20 μL. Cycling conditions were 95^◦^C 3 min, 32 cycles of 95^◦^C 1 min, 58^◦^C 1 min, 72^◦^C 1 min, followed by 72^◦^C for 5 min. PCR products were run on the Applied Biosystems 3730xl DNA Analyzer using GS500LIZ internal size standard and analyzed using GeneMapper v5. Repeat size of the modal allele was inferred based on the size of the PCR product in base-pairs. Expansion indices were calculated from GeneMapper peak height data as described^32^ using a 5% relative peak height threshold. These were calculated relative to the modal allele of the more stable striatal peak, which was typically the lowest modal allele length amongst all tissues at 35 weeks and assumed to be the closest approximation to the inherited repeat length.

### Western Blot

Cerebellum, brain stem, and cerebral cortex, was collected from *Atxn1^2Q/2Q^*, *f-ATXN1^146Q/2Q^* and *Atxn1^175Q/2Q^* mice at 2 weeks of age. Samples were homogenized using a tissue grinder in 500 ul of Tris Triton lysis buffer (50mM Tris, pH 7.5, 100mM NaCl, 2.5mM MgCl2, 0.5% Triton X-100) that included MilliporeSigma protease inhibitors II and III and a Roche Complete Mini Protease inhibitor tablet. Homogenized samples were shaken at 1500 rpm at 4°C for 1 hour, frozen and thawed in liquid nitrogen and 37°C water bath 3 times, and centrifuged at 21,000xg for 10 min at 4°C. Samples containing 30μg total protein were boiled in Laemmli loading buffer and run on a 4%–20% Bio-Rad precast gel. Protein was transferred to a nitrocellulose membrane using the BioRad Trans-Blot Turbo system. Blots were cut at approximately 75kDa and blocked overnight at 4°C in 5% milk PBST (phosphate-buffered saline, 0.1% Tween 20). Blots were probed overnight at 4°C 1:2500 with the ATXN1 antibody 11750^33^ or 1:10,000 with α-Tubulin antibody (MilliporeSigma T5168) diluted in 5% milk PBST. Blots were washed 3 times with PBST. ATXN1 blots were then placed in 5% milk PBST plus 1:2500 rabbit specific HRP antibodies (GE Healthcare) while α-Tubulin blots were placed in 5% milk PBST plus 1:10,000 mouse specific HRP antibodies (GE Healthcare) at room temperature for 4 hours. Blots were washed 3 times with PBST and then ATXN1 blots were washed with Super Signal West Dura (Thermo Fisher Scientific) while α-Tubulin blots were washed with Super Signal West Pico (Thermo Fisher Scientific) detection reagents. Blots were imaged on an ImageQuant LAS 4000. For the DARPP-32 western, striatal tissue was homogenized in sample buffer as above, blotted and probed with rat monoclonal anti-DARPP-32 (R&D Systems, MAB4230) at 1:1000 and anti-rat IgG HRP secondary at 1:2500 (GE Healthcare, NA935).

### RT-qPCR

One half of each brain region or tissue from each mouse was homogenized in 500 mL TRIzol Reagent (Thermo Fisher Scientific, 15596026). RNA isolation was done per the manufacturer’s instructions. cDNA was synthesized in duplicate using 500 ng RNA in 10 uL iScript Advanced cDNA Synthesis Kit (Bio-Rad, 172-5038). Reactions were diluted 1:5 in water. qPCR was done using 2 uL diluted cDNA in 10 uL Roche Probes Master (04707494001) reactions on a Roche 480 Lightcycler. Target gene and reference gene reactions were amplified in separate wells under cycling conditions of 95C for 10 s, 60C for 10 s for 35 cycles. Primers used, ATXN1 Forward (5’AGAGATAAGCAACGACCTGAAGA) and ATXN1 Reverse (5’CCAAAACTTCAACGCTGACC) with Roche probe 67 (04688660001). NMJ qPCR was done using 2 ul diluted cDNA in 10 ul Roche Sybr Green (04707516001) reactions under cycling conditions of 95C for 10 s, 50C for 10 s and 72C for 10 s for 45 cycles. Primers used, Chrna1 Forward (5’ CATCGAGGGCGTGAAGTACA) and Chrna1 Reverse (5’ ATTCCTCAGCGGCGTTATTG) and Musk Forward (5’ TGAGAACTGCCCCTTGGAACT) and Musk Reverse (5’ GGGTCTATCAGCAGGCAGCTT). Reference genes from Roche (Proprietary sequence) mGapdh (5046211001) and mACTB (05046190001). Cq (quantitation cycle) values were determined using the Roche second derivative maximum calculation. Relative quantification was done using standard 2^DD^Cq.

### Total mouse/human RT-qPCR assay

Total *ATXN1* RNA levels were measured using primers and probes that measures both human and mouse RNA levels in the same reaction. The assay was designed with a forward primer that binds to Exon 6 of the mouse sequence which is common for all genotypes. (mAtxn1 Forward 5’AAGAAAGACACCACCAGAACC) The reverse primer was designed to bind to an identical sequence shared by mouse exon 7 and human exon 8. (m+hATXN1 Reverse 5’GATTTCTGTAGGGGATCCAGGC) This will generate two highly similar but unique amplicons using a single set of primers. Quantitation of the two amplicons will be achieved by utilizing two probes. Probes were designed to bind to unique sequences within the amplicons conferring specificity for human or mouse sequence: mAtxn1 Ex6-Ex7 (56-FAM/CCACT GCCA/ZEN/GCCTAAAGAACCCA/3IABkFQ) or hAtxn1 Ex8 (5HEX/CCAGAGCTG/ZEN/C TGTTGGCGGATTGTA /3IABkFQ) qPCR was optimized for primer concentration, probe concentration and checked for reaction efficiency. Efficiency for mouse reaction 1.926, human 1.925.

### CT Scan

Mice were anesthetized with 2.5% isoflurane and CT scans were performed using a Sophie G8 uPET/uCT imaging system (PerkinElmer). The x-ray source for the CT scans were set to 59 kVp and 100uA and the resulting voxel size was 200um isotropic. 3-dimensional CT images were analyzed using Imaris 9.8 (Oxford Instruments). Kyphosis index was determined based on where the distance is calculated from a horizontal line drawn from the center of the C7 vertebrae to the center of the pelvis, and a vertical line from the apex spine curvature to the intersection of the horizontal line.^20^ Measurement points were manually added in 3-dimensional space using the Imaris spots module.

### Behavior Methods

A behavioral battery was used to assess neuromotor and neurocognitive function in individual animals over time. Testing was repeated at 6, 12, 18, 26, 30, and/or 31 weeks of age.

### Rotarod

A rotarod apparatus (Ugo Basile) was used to assess motor coordination and balance. Testing occurred over four consecutive days and each test day consisted of 4 individual trials (∼15 min intertrial interval) where mice were placed on a rotating rod that accelerated from 5-50 RPM in 1 RPM steps over the course of a 5 min interval (∼7 sec ramp). The trial ended when mice fell from the rod, when they failed to continuously walk on the rod (held on to the rod instead of walking for two full 360° rotations), or after the 5 min maximum trial time was reached. The apparatus was cleaned with 70% ethanol between animals and between each trial. The latency to fall (sec) was averaged across the 4 trials in individual animals for analysis.

### Barnes maze

A Barnes maze apparatus (San Diego Instruments) was used to assess spatial learning and memory.^34^ The apparatus consisted of a white circular ABS plastic arena (36” diameter, 36” high) with 20 2” holes evenly spaced along the perimeter. One of the holes contained a recessed target box that allowed the mice an escape from the open space of the arena. Contextual markers were located on the walls around the maze to provide spatial cues and the apparatus was brightly illuminated (∼250 lux) during testing. Training consisted of 4 trials a day (∼15 min intertrial interval) for 4 days and trial began by placing the mouse in the center of the arena and ended when the mouse entered the goal box or after the 3 min maximum test time had elapsed. The location of the goal box remained the same across training trials. A probe test was conducted the following day where the target hole was blocked off and mice were allowed to explore the arena during a single 90 sec trial. The apparatus was cleaned with 70% ethanol between animals and between each trial. The latency (sec) to enter the escape hole was averaged across the 4 trials in individual animals each day during training and the time spent (sec) in the goal versus other 3 quadrants of the maze (+1, opposite, -1) was assessed for the probe test.

### Open Field

An open field arena was used to assess exploratory locomotor activity and consisted of a white rectangular box (20”W x 20”L x 10” H) illuminated by overhead LED lights (∼150 lux). Mice were allowed to explore the open field arena for 30 min and ANY-maze video tracking software was used to measure the total distance traveled (m) and average speed (m/sec) in individual animals. The apparatus was cleaned with 70% ethanol between animals.

## Muscle Strength Assays

### In Vivo Torque

Mice were anesthetized with isoflurane and maximal isometric torque of the anterior crural muscles was measured. Sterilized platinum needle electrodes were placed through the skin near the left common peroneal nerve and connected to a stimulator (Models S48, Grass Technologies, West Warwick, RI) and stimulus isolation unit (SIU5, Grass Technologies, West Warwick, RI). The contractile function of the anterior crural muscles was assessed by measuring isometric torque (variable voltage (3-10V), 150-ms train, and 0.1-ms pulses) every minute until peak torque was achieved.

### Ex vivo muscle preparation

EDL force production was assessed according to methods described previously.^35^ Briefly, mice were anesthetized with sodium pentobarbital (75–100 mg/kg body mass) and EDL muscles were excised and mounted in a 1.2 mL bath assembly with oxygenated (95:5% O2/CO2) Krebs Ringer bicarbonate (Krebs) buffer maintained at 25°C. Muscles were adjusted to their anatomical optimal length (Lo) based on resting tension, with length being measured from the distal myotendinous junction to the proximal myotendinous junction. Muscles were equilibrated in the bath for 10 minutes before performing isometric contractions using maximal voltage (150 V) for 200 ms at 175 Hz every 2 mins until peak isometric force was achieved.

### Immunofluorescence

TA muscles from each mouse line were prepared in melting isopentane for 30s and 10 µm transverse cryosections were obtained (Leica CM3050 S). Sections were fixed in acetone at -20°C for 15 min and subsequently washed 3x in PBS before being blocked in 5% goat serum for 30 mins at RT. Sections were then incubated for >1h in primary antibody (rat monoclonal anti-Laminin 1:500; Sigma, L06631) at RT, washed 3x with PBS, and incubated with Alexa Fluor 488 Goat anti-Rat IgG (ThermoFisher, A-11006) secondary (1:1000) for 30 mins at RT. Finally, sections were washed 3x in PBS and mounted in ProLong Gold Antifade with 4′,6-diamidino-2-phenylindole (DAPI) to visualize nuclei (ThermoFisher Scientific). Images were acquired on a Leica DM5500 B microscope equipped with a Leica HC PLAN APO 10× objective and stitched together with LASX software (Leica) to allow visualization of the entire TA. SMASH – semi-automatic muscle analysis using segmentation of histology software was used to analyze and quantify centrally located nuclei, fiber number, and fiber size.^36^

## References

1. Orr, H.T. et al. Expansion of an unstable trinucleotide CAG repeat in spinocerebellar ataxia type 1. Nat. Genet. 4, 221–226 (1993).

2. Hannan, A.J. Tandem repeats mediating genetic plasticity in health and disease. Nat. Rev. Genetics 19, 286–298 (2018).

3. Robitaille, Y., Schut, L. & Kish, S.J. Structural and immunocytochemical features of olivopontocerebellar atrophy caused by the spinocerebellar ataxia type1 (SCA-1) mutation define a unique phenotype. Acta Neuropath. 90, 572–581 (1995).

4. Schöls, L. et al. Spinocerebellar ataxia type 1: clinical and neurophysiological characteristics in German kindreds. Acta Neurol. Scand. 92, 478–485 (1995).

5. Bürk, K. et al. Autosomal dominant cerebellar ataxia type 1 clinical features and MRI in families with SCA1, SCA2, and SCA3. Brain 119, 1497–1505 (1996).

6. Bürk, K. et al. Cognitive deficits in spinocerebellar ataxia type 1, 2, and 3. J. Neurol. 250, 207–211 (2003).

7. Rüb, U. et al. Spinocerebellar ataxia type 1 (SCA1): new pathoanatomical and clinic-pathological insights. Neuropath. Appl. Neurobio. 38, 665–680 (2012).

8. Watase, K. et al. A long CAG repeat in the mouse Sca1 locus replicates SCA1 features and reveals the impact of protein solubility on selective neurodegeneration. Neuron 34, 905–919 (2002).

9. Hayashi, S., Lewis, P., Pevny, L. & McMahon, A.P. Efficient gene modulation in mouse epiblast using a *Sox2Cre* transgenic mouse strain. Mech. Develop. 119S, S97–S101 (2002).

10. Tronche, F. et al. Disruption of the glucocorticoid receptor gene in the nervous system results in reduced anxiety. Nat Genet. 23, 99–103 (1999).

11. Lalonde, R. & Strazielle, C. Brain regions and genes affecting limb-clasping responses. Brain Res. Rev. 67, 252–259 (2011).

12. Keiser, M.S., Boudreau, R.L. & Davidson, B.L. Broad therapeutic benefit after RNAi expression vector delivery to deep cerebellar nuclei: Implications for spinocerebellar ataxia type 1 therapy. *Mol*. Therapy 22, 588–595 (2014).

13. Nitschke, L. et al. miR760 regulates ATXN1 levels via interaction with its 5’ untranslated region. Genes & Devel. 34, 1147–1160 (2020).

14. Reetz, K., et al. Genotype-specific patterns of atrophy progression are more sensitive than clinical decline in SCA1, SCA3 and SCA6. Brain 136, 905–917 (2013).

15. Koscik, T.R. et al. Brainstem and striatal volume changes are detectable in under 1 year and predict motor decline in spinocerebellar ataxia type 1. Brain Commun. 2, 1–13 (2020).

16. Dang, M.T. et al. Disrupted motor learning and long-term synaptic plasticity in mice lacking NMDAR1 in the striatum. Proc. Natl. Acad. Sci. USA 103,15254–15259 (2006).

17. Wang, N. et al. Neuronal targets for reducing mutant huntingtin expression to ameliorate disease in a mouse model of Huntington’s disease. Nat Med. 20, 536–542 (2014).

18. Watase, K., Venken, K.J.T, Sun, Y., Orr, H.T. & Zoghbi, H.Y. Regional differences of somatic CAG repeat instability do not account for selective neuronal vulnerability in a knock-in mouse model of SCA1. Hum. Mol. Genet, 12, 2789–2795 (2003).

19. Pinto, R.M. et al. Patterns of CAG repeat instability in the central nervous system and periphery in Huntington’s disease and in spinocerebellar ataxia type 1. Hum. Mol. Genet. 29, 2551–2567 (2020).

20. Laws, N. & Hoey, A. Progression of kyphosis in *mdx* mice. J. Appl. Physiol. 97, 1970–1977 (2004).

21. Miniou, P., Tiziano, D., Frugier, T., Roblot, N., Le Meur, M. & Melki, J. Gene targeting restricted to mouse striated muscle lineage. Nuc. Acids Res. 27, e27.

22. Pratt, S.J.P., Shah, S.B., Ward, C.W., Kerr, J.P., Stains, J.P., & Lovering, R.M. Recovery of altered neuromuscular junction morphology and muscle function in mdx mice after injury. Cell. Mol. Life Sci. 72, 153–164 (2015).

23. Handler, H.P., et al. Disrupting ATXN1 Nuclear Localization in a Knock-in SCA1 Mouse Model Improves a Spectrum of SCA1-Like Phenotypes and their Brain Region Associated Transcriptomic Profiles. Neuron, in press (2023).

24. Gall-Duncan, T., et al. Antagonistic roles of canonical and alternative RPA in tandem CAG repeat diseases. bioRxiv, doi: https://doi.org/10.1101/2022.10.24.513561 (2022)

25. Boucher, C. and Johnson, K. Triplet repeats in neuromuscular disorders. Ann. Med. 27, 3–5 (1995).

26. Abe, K., Kameya, T., Tobita, M., Konno, H. and Itoyama, T. Molecular and clinical analysis on muscle wasting in patients with spinocerebellar ataxia type 1. Muscle & Nerve, 19, 900–902 (1996).

27. Fukazawa, S.H. et al. Clinical features and natural history of spinocerebellar ataxia type 1. Acta Neurol. Scand. 93, 64–71 (1996).

28. Genis, D. et al. Clinical, neuropathologic, and genetic studies of a large spinocerebellar ataxia type 1 (SCA1) kindred: (CAG)n expansion and early premonitory signs and symptoms. Neurology 45, 24–30 (1995).

29. Orengo, J.P. et al. Motor neuron degeneration correlates with respiratory dysfunction in SCA1. Dis. Model. Mech. 11, 1–7 (2018).

30. Ran, F.A., Hsu, P.D., Wright, J., Agarwala, V., Scott, D.A. &s Zhang, F. Genome engineering using the CRISPR-Cas9 system. Nat. Protocols 8, 2281–2308 (2013).

31. Gamache, J., et al. Factors other than hTau overexpression that contribute to taupathy-like phenotype in rTg4510 mice. Nat. Comm. 10, 2479 (2019).

32. Lee, J.M., et al. A novel approach to investigate tissue-specific trinucleotide repeat instability. BMC Syst. Biol. 4, 29. Doi: 10.1186/1752-0509-4-29.

33. Servadio, A., Koshy, B., Armstrong, D., Antalffy, B., Orr, H.T. & Zoghbi, H.Y. Expression analysis of the ataxin-1 protein in tissues from normal and spinocerebellar ataxia type 1 individuals. Nat. Genet. 10: 94–98 (1995).

34. Illouz, T. et al. Unbiased classification of spatial strategies in the Barnes maze. Bioinformatics 32, 3314–3320 (2016).

35. Moran, A. L., Warren, G. L. & Lowe, D. A. Soleus and EDL muscle contractility across the lifespan of female C57BL/6 mice. Exp. Gerontol. 40, 966–975 (2005)

36. Smith, L.R. & Barton, E.R. SMASH – semi-automatic muscle analysis using segmentation of histology: a MATLAB application. Skeletal Muscle 4:21 (2014).

